# In search of a unifying theory of white matter aging: improving the understanding of tract-wise degeneration using multi-parametric signatures of morphometry and microstructure

**DOI:** 10.1101/2023.03.14.532658

**Authors:** Tyler D. Robinson, Yutong L. Sun, Paul T. H. Chang, J. Jean Chen

## Abstract

While tract-wise differences in volume and microstructure are common targets of investigation in age-related changes in the white matter (WM), there has been relatively little exploration into other attributes of tract morphometry or its relation to microstructure in vivo, and limited understanding on how they jointly inform the interpretation of the WM aging trajectory. This study examines ten WM tracts for tract-wise differences in morphometry (i.e. volume, length, and volume-to-length ratio) and microstructural integrity (i.e. fractional anisotropy, mean diffusivity, axial diffusivity, and radial diffusivity) using diffusion MRI data from the Human Connectome Project in Aging (HCP-A) with the goal of laying the foundation for a unified model of age-related WM microstructure-morphometry trajectories with a special focus on sex differences. Results indicated widely heterogeneous patterns of decline and morphometry-microstructural associations across tracts. Multi-parametric signatures of decline suggest stages or mechanisms of degeneration that differ between sexes. This work highlights the value of integrating microstructural and morphometric measures of WM health instead of observing them separately, suggesting multiple modes of WM degeneration.

## INTRODUCTION

Changes in function, structure, and microstructure in the human brain occur throughout the adult lifespan (Baker et al., 2014; Bartzokis, 2004; Bookheimer et al., 2019; Buyanova & Arsalidou, 2021; Chad et al., 2018; Kiely et al., 2022; Kodiweera et al., 2016; Piguet et al., 2009). Earlier prediction of regional declines is invaluable to medical intervention for the postponement or reversal of progressive cognitive and structural decline (de Groot et al., 2015). To this end, evidence suggests that early microstructural degeneration can be observed in the white matter (WM) prior to their emergence in cortical regions (Debette & Markus, 2010; Hu et al., 2021; Madden et al., 2012; Seiler et al., 2018). Consistent changes in white matter macro-/microstructure and histology have been demonstrated in advancing age (Baker et al., 2014; Bastin et al., 2010; Choy et al., 2020; Ouyang et al., 2021; Tang et al., 1997), with overall volume decreases identified in normal aging (Bastin et al., 2010; Piguet et al., 2009; Tang et al., 1997). Notably, WM volume decreases by up to 28% in adult aging (Pakkenberg & Gundersen, 1997), but there is still a limited understanding of the microstructural variations underlying this volume reduction. In order to understand the mechanisms underlying WM decline, it is important to incorporate knowledge of age-related changes in WM morphometry and microstructure (Bartzokis, 2004; Benitez et al., 2018; Maffei et al., 2021; Yendiki et al., 2011). In turn, examining the relationship between this interaction and the WM developmental trajectory can help further elucidate the processes that govern WM degeneration in aging. Building a more comprehensive modeling of age-related WM changes offers a tool to detect degeneration at earlier stages.

While WM volume loss in aging has been established, the regional distribution of this volume loss is as of yet under-defined. Decreases in cumulative fiber length (i.e. the total length across the brain as extrapolated from histological samples) have been observed in both humans and animals using diffusion MRI and microscopy, respectively (Baker et al., 2017; Marner et al., 2003). This loss can in part be attributed to the preferential culling of vulnerable axons, as fibers with lower axon diameter and less dense myelin sheaths are notably more vulnerable to this process (Tang et al., 1997). This further suggests that a significant contributor to WM morphometrical losses in aging may be a reduction in the total number of fibers and a concomitant increase in mean axon diameter in surviving tissue, as opposed to changes in the length of surviving axons. As WM fibers with higher diameter are also typically longer (Liewald et al., 2014), aging tracts should demonstrate increases in the length-to-volume ratio as shorter, narrower fibers are culled and the longer surviving fibers comprise a greater portion of the WM tissue. However, to what degree this potential effect contributes to changes in WM morphometry, considering the general atrophy of brain volume with age (Azevedo et al., 2019; Fjell et al., 2009, 2014; Sala et al., 2012), is unclear. Individual fiber tracts in the white matter exhibit varying degrees of vulnerability to age-related fiber culling (Choy et al., 2020), potentially giving rise to previous observations of both decreasing and increasing mean fiber length in specific major white matter bundles including the splenium and superior longitudinal fasciculus (Ouyang et al., 2021), calling into question the assumption that overall decreases in fiber length correspond to regional decreases in tract length. Moreover, the tools with which living WM can be accurately delineated into discrete tracts are both relatively new and continually improving (Maffei et al., 2021). Alternate mechanisms of shortening in measurable fibre length, such as damage and delayed clearing of remaining debris (Avellino et al., 1995) or the presence of early-stage degenerative conditions targeting short-range structural connections (Wu et al., 2022) may also influence measures of mean tract length. Morphometrical research has noted cross-sectional tract-wise differences in age-related decline with regards to volume and microstructure (Bastin et al., 2010; Choy et al., 2020), but in the context of aging, it is also important to assess normalized fibre length in order to account for sex differences and age-related general atrophy, thereby gleaning the proportional instead of absolute changes in different WM tracts (Bajada et al., 2019). As many existing works do not address potential sex differences, distinctions between sex-specific trajectories of decline remain unclear.

Reported to show alterations earlier than cortical morphometry (Debette & Markus, 2010; Hu et al., 2021; Madden et al., 2012; Seiler et al., 2018), diffusion MRI metrics such as fractional anisotropy (FA) and mean diffusivity (MD) offer proxy measures for WM microstructural integrity (Madden et al., 2012; Stadlbauer et al., 2008). Phenomena such as culling of axonal fibers and demyelination of surviving axons results in measurable changes in diffusion, such that mean diffusivity increases and fractional anisotropy decreases as axon bundles become less dense (Choy et al., 2020; Fan et al., 2019; Kodiweera et al., 2016). Existing research has also identified heterogeneous rates of decline in different WM regions, such as sections of the corpus callosum, and broader anterior and posterior regions of the brain (C. L. Armstrong et al., 2004; Brickman et al., 2012; Fan et al., 2019; Kiely et al., 2022; Seiler et al., 2018). Declines in microstructural integrity in aging WM are associated with both WM hyperintensities and implicated in declining cognitive performance (Debette & Markus, 2010; Hu et al., 2021). This knowledge is of considerable interest for anticipating pathological changes and prioritizing interventions (Debette & Markus, 2010). Tract-wise studies and inter-tract comparisons of WM microstructural variations in aging remain limited. Recent work by Schilling et al. involved the largest data set and number of tracts to date (Schilling et al., 2022), and highlighted the heterogeneity in multiple morphometric dimensions across the WM, which is a strong motivation for this study.

We reasonably expect microstructural variations to underlie morphometrical variations in aging (N. M. Armstrong et al., 2020; Hoagey et al., 2019; Taki et al., 2011), but interaction between morphometrical and microstructural declines represents a sparsely addressed topic (Bastin et al., 2010; Kezele et al., 2008; Seiler et al., 2018), as most studies compare microstructural and morphometric age variations without associating them (Sala et al., 2012; Schilling et al., 2022). Prior investigation notes a decrease in overall fiber count with age that could account for measurable changes in WM volume (Marner et al., 2003). This would pose microstructural declines as a direct cause of morphometrical changes in volume. However, histological findings also note that surviving WM fibers are thicker with advancing age (Fan et al., 2019, 2020). Moreover, while age may result in lower density in fiber bundles (Choy et al., 2020; Fan et al., 2019; Kiely et al., 2022; Marner et al., 2003), changes in density may not necessitate a uniform decreases in volume, as space within fiber bundles is occupied by intercellular fluid (Kodiweera et al., 2016). Uncertainty in these processes make it difficult to anticipate the relationships between microstructural decline and changing WM morphometry with age. This study sought to map changes in WM morphometry and microstructure, examine their interactions, and compare those changes across individual WM tracts. Tract-wise FA and MD, as well as axial diffusivity (AD) and radial diffusivity (RD) are used as the target microstructural metrics of this study. The morphometry of each tract is quantified in measures of tract-wise mean length, volume, and volume-length ratio, and these are, in turn, compared to the microstructural measures. We hypothesize that: (1) due to the potential causality between microstructural and morphometric variations, age-related microstructural differences will be greater in tracts that demonstrate greater age-related morphometrical differences, and (2) given patterns of preferential WM fiber culling (Marner et al., 2003; Tang et al., 1997), and associations between axon diameter and axon length (Liewald et al., 2014), tracts that become thinner with age will also have higher mean tract lengths.

## METHODS

### Subjects and data acquisition

This study involved data from five hundred and thirty-five healthy adult subjects (300 female, aged 36-100 years) of the Human Connectome Project in Aging (HCP-A) dataset (OMB Control# 0925-0667) (Bookheimer et al., 2019; Harms et al., 2018). Of this sample, thirty-two subjects were excluded due to MRI artifacts resulting in notable errors when producing WM masks. An additional three subjects (100 years of age) were excluded as outliers. This resulted in a final sample size of 500 subjects (294 female, aged 36-89). All subjects were in good health and without pathological cognitive impairment (i.e. stroke, clinical dementia).

Each data set included a whole-WM T1-weighted structural MRI (0.8 mm isotropic resolution) and diffusion-weighted MRI (dMRI), collected on one of four matched Siemens Prisma 3T MRI scanners, with a 1.5 mm isotropic voxel resolution, MB=4, with 93 directions at b=1500s/mm^2^. Further parameters for structural and diffusion image acquisition can be found in (Harms et al., 2018).

### Image analysis

dMRI data were corrected for eddy-current and susceptibility-related distortions via EDDY. Diffusion data were used to identify and reconstruct eighteen major white matter tracts using the Tracts Constrained by Underlying Anatomy (TRACULA) tool in Freesurfer version 7.2.0 (Maffei et al., 2021; Yendiki et al., 2011). FMRIB Software Library version 6.0.5 (FSL)’s fslmaths function was then used to produce per subject masks of each WM tract in local space. A 99^th^ percentile voxelwise apparent diffusion coefficient (ADC) intensity threshold was applied to the generated tracts to exclude potential outliers, followed by a 20% threshold on the resulting masks to generate the final masked regions of interest (ROI) used throughout this study. The eighteen reconstructed tracts were collapsed bilaterally to produce ten tracts of interest: The major forceps (Fmajor), minor forceps (Fminor), anterior thalamic radiation (ATR), Cingulum Angular Bundle (CAB), Cingulate Gyrus (CCG), Corticospinal Tract (CST), Inferior Longitudinal Fasciculus (ILF), Superior Longitudinal Fasciculus Parietal (SLFP), Superior Longitudinal Fasciculus Temporal (SLFT), and the Uncinate Fasciculus (UNC) (**Figure 1**). In single tract analyses, Rosner’s tests were performed to identify and exclude volume outliers per tract. The resulting subject counts were as follows: Fmajor (N=493), Fminor (N=493), ATR (N=497), CAB (N=490), CCG (N=495), CST (N=491), ILF (N=497), SLFP (N=497), SLFT (N=495), and UNC (N=495).

**Figure 1:**
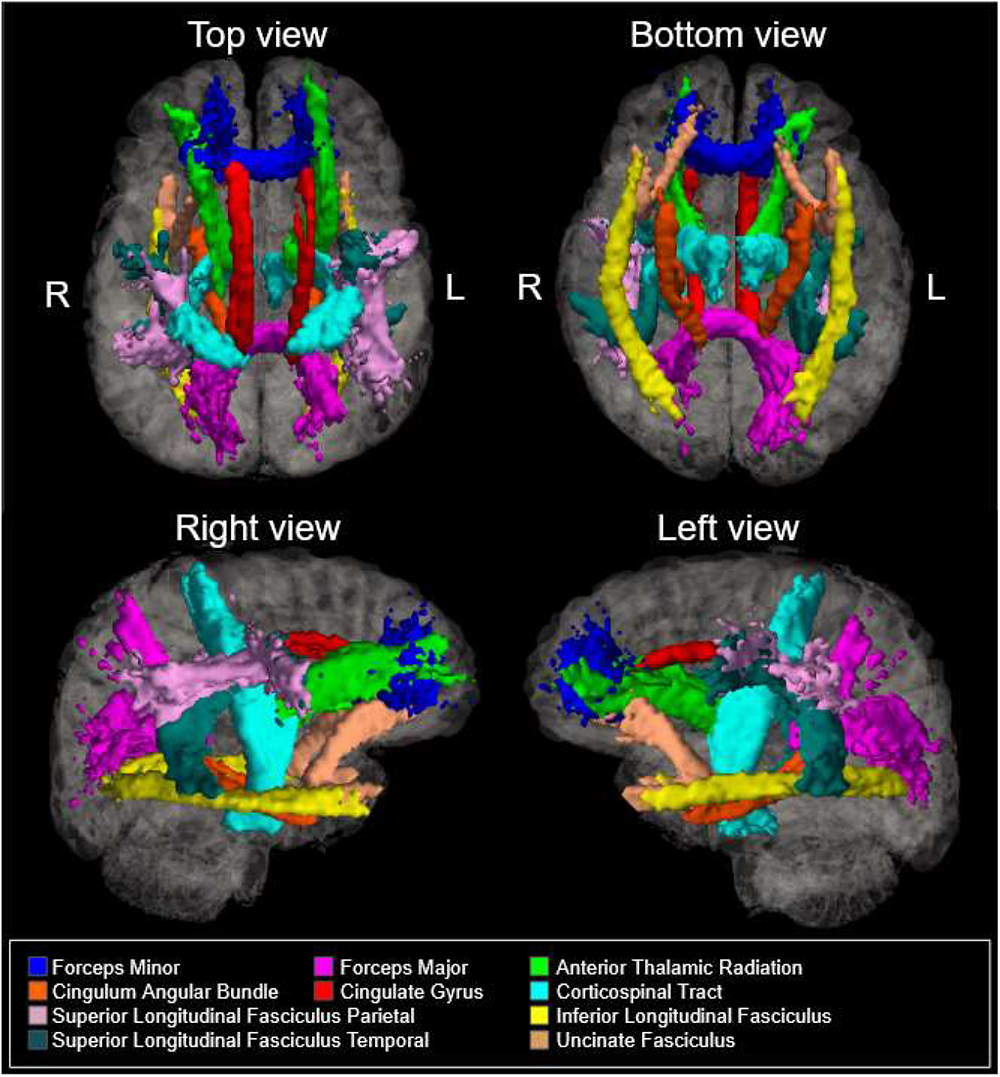
Ten bilateral tracts of interest reconstructed Freesurfer’s TRACULA package

#### Morphometry

From these segmented regions, morphometry measures were calculated for tract volume, length, and volume-to-length ratio (VLR) using Python (Python Software Foundation) along with fslmaths (FMRIB Software Library version 6.0.5 (FSL)):

1. Tract volume:

a. Raw tract volume (“Volume”): calculated as the total number of voxels (2x2x2mm^3^) included in each masked tract region.
b. Normalized tract volume (“Volume_norm_”): calculated as tract volume divided by subject-specific intracranial volumes (ICV). This represents the tract volume as a percentage of the brain volume, thereby reducing the effect of inherent inter-subject variability in tissue volumes.
c. Percent tract volume change (“Volume_perc_”): calculated as individual raw tract volume divided by the mean raw tract volume of the youngest 10% of subjects.
2. Tract length

a. Raw tract length (“Length”): calculated as the mean streamline length from 1500 streamlines per tract in TRACULA.
b. Normalized tract length (“Length_norm_”): the raw tract length divided by the length of an subject-specific length of age-stable white matter tract (the cingulate gyrus (CCG)), as identified from its relatively stable raw tract length identified in the raw values. This reduces the effect of inherent inter-subject variability in tract lengths.
c. Percent tract length change (“Length_perc_”): calculated as individual raw tract length divided by the mean raw tract length of the youngest 10% of subjects.
3. Tract volume-to-length ratio (VLR)

a. Raw VLR (“VLR”): calculated as the ratio of tract volume to length. This represents the “thinness” of a tract relative to a reference tissue.
b. Normalized VLR (“VLR_norm_”): calculated as the ratio of ICV-corrected tract volume to CCG-corrected tract length. This represents the “thinness” of a tract relative to a reference tissue.
c. Percent tract VLR change (“VLR_perc_”): calculated as individual raw tract VLR divided by the mean raw tract VLR of the youngest 10% of subjects.

#### Microstructure

Fractional anisotropy (FA), mean diffusivity MD, axial diffusivity (AD), and radial diffusivity (RD) maps were derived using Dipy’s DKI tool, to provide kurtosis-corrected DTI metrics (given the high b-value used in the HCP-A dMRI acquisitions). Tract masks generated in the morphometric assessment were applied to the local space diffusion images using fslmaths to generate mean FA and mean diffusivity values for each tract of interest of each participant. As with morphometric measures, baseline normalized values (“FA_perc_”, “MD_perc_”, “AD_perc_”, “RD_perc_”) were calculated for each microstructural measure to determine percent difference from the youngest 10% of the sample.

### Statistical analyses

#### Morphometry

In these analyses we assessed the relationships between morphometry, age and sex using multivariate linear and quadratic regression. All analyses in this study were conducted in R (version 4.1.1). First, between-subjects ANOVA was used to determine the presence of sex differences in tract specific Length_norm_, Volume_norm_, and VLR_norm_. Age was then regressed onto tract-wise Length_norm_, Volume_norm_, and VLR_norm_ with sex as a covariate of no interest. Subsequent assessment of sex differences in tract-wise morphometry was conducted via sequential multivariate regressions of sex onto tract Length_norm_, Volume_norm_, and VLR_norm_ using age as a covariate of no interest. False detection rate (FDR) adjustment was applied to all resulting p-values to correct for multiple comparisons.

#### Microstructure

Age-related microstructure differences were assessed via multivariate regression by first regressing age onto tract-wise MD_perc_ and mean FA_perc_ values derived from voxel-count weighted tract MD_perc_ and FA_perc_ for each of the ten tracts using sex as a covariate of no interest. Subsequent assessment of sex differences in microstructure was conducted via sequential multivariate regressions of sex on to mean MD_perc_ and mean FA_perc_, using age as a covariate of no interest. FDR adjustment was again applied to the resulting p-values.

#### Morphometry vs. microstructure

Two multivariate regression models were constructed to assess the relationships between morphometry, microstructure, age, and sex in our sample. These models were first applied to whole-WM Length_perc_ and VLR_perc_ values, then to the individual values for each tract of interest:

● *Model 1. microstructure vs. morphometry across ages:* assesses the degree to which morphometric metrics account for the age-related variability in mean normalized tract length, volume, and VLR, with subject sex as a covariate of no interest. That is,

○ {FA_perc_, MD_perc_} = f(Length_perc_+Sex)
○ {FA_perc_, MD_perc_} = f(VLR_perc_+Sex)
● *Model 2. microstructure vs. morphometry between sexes:* assess the degree to which morphometric metrics account for sex-related differences in normalized tract length, volume and VLR, with subject age as a covariate of no interest. That is,

○ {FA_perc_, MD_perc_} = f(Length_perc_+Age)
○ {FA_perc_, MD_perc_} = f(VLR_perc_+Age)
○ One set of models is produced for each sex.

#### Canonical Correlation Analysis

As a method for multimodal fusion, canonical correlation analysis (CCA) was conducted to examine the degrees to which our variables of interest contributed to age-related differences in morphometry and microstructure respectively. Two column vectors were computed using our morphometric (*V*_1_) variables (Length_perc_, Volume_perc_) and microstructural (*U*_1_) variables (AD_perc_, RD_perc_, FA_perc_) in Matlab. The linear combinations of *V* and *U* vectors which produced the highest correlation were then calculated to determine on a tract-wise basis which variables loaded most strongly to age differences by tract. This and following analyses substituted AD and RD variables for the MD variable used in previous analyses as examining the axes of diffusion separately was necessary to clarify relationships between FA and other diffusion metrics as they relate to microstructural deterioration.

#### Cross-Correlation Analysis

Cross-correlations functions (CCF) were calculated in Matlab (Mathworks Inc., Natick, USA) to determine the temporal relationships between morphometry and microstructure variations, as well as between those in measures within each category. To reduce ambiguity in the CCF interpretation, only variables demonstrating significant non-linear age effects on a tract-wise basis were examined in this analysis, using a curvature threshold of 0.0005, where curvature was computed using Matlab’s *curvature* function. CCFs were calculated based on the models of age effects on six paired tract-wise microstructure/morphometry measures, along with cross-comparisons for AD_perc_/RD_perc_ vs FA_perc_ and Length_perc_ vs Volume_perc_:

● Length/Microstructure: Length_perc_ vs AD_perc_, Length_perc_ vs RD_perc_, Length_perc_ vs FA_perc_
● Volume/Microstructure: Volume_perc_ vs AD_perc_, Volume_perc_ vs RD_perc_, Volume_perc_ vs FA_perc_
● AD_perc_ vs FA_perc_
● RD_perc_ vs FA_perc_
● Length_perc_ vs Volume_perc_

## 3 RESULTS

### Morphometry

#### Age Differences

Initial analysis of relationships between age and raw morphometric measures identified significantly negative associations between age and raw tract Length in Fminor, CAB, CST, ILF, SLFP, SLFT, and UNC, along with significant negative associations between age and raw tract Volume in Fmajor, Fminor, ATR, CAB, CCG, ILF, SLFP, SLFT, and UNC, and between age and raw tract VLR in Fmajor, Fminor, ATR, CAB, CCG, ILF, SLFP, SLFT, and UNC. Echoing the raw morphometric analyses, negative relationships between age and tract Length_norm_ were identified in Fminor, CAB, CST, ILF, SLFP, SLFT, and UNC. Negative relationships between age and tract Volume_norm_ and VLR_norm_ were identified in Fmajor, Fminor, ATR, CAB, CCG, ILF, SLFP, SLFT, and UNC, as well as between age and tract VLR_norm_ in the Fmajor, the Fminor, ATR, CAB, CCG, and UNC (**Figure 2**). Following baseline normalization, quadratic, rather than linear regression was found to provide a better fit for age effects on Length_perc_ in the Fminor, ATR, CAB, CCG, ILF, and SLFP. Similarly, quadratic, rather than linear declines provided better fits for age effects on Volume_perc_ in the CCG, ILF, and SLFP (**Supplemental Figure 1**).

**Figure 2:**
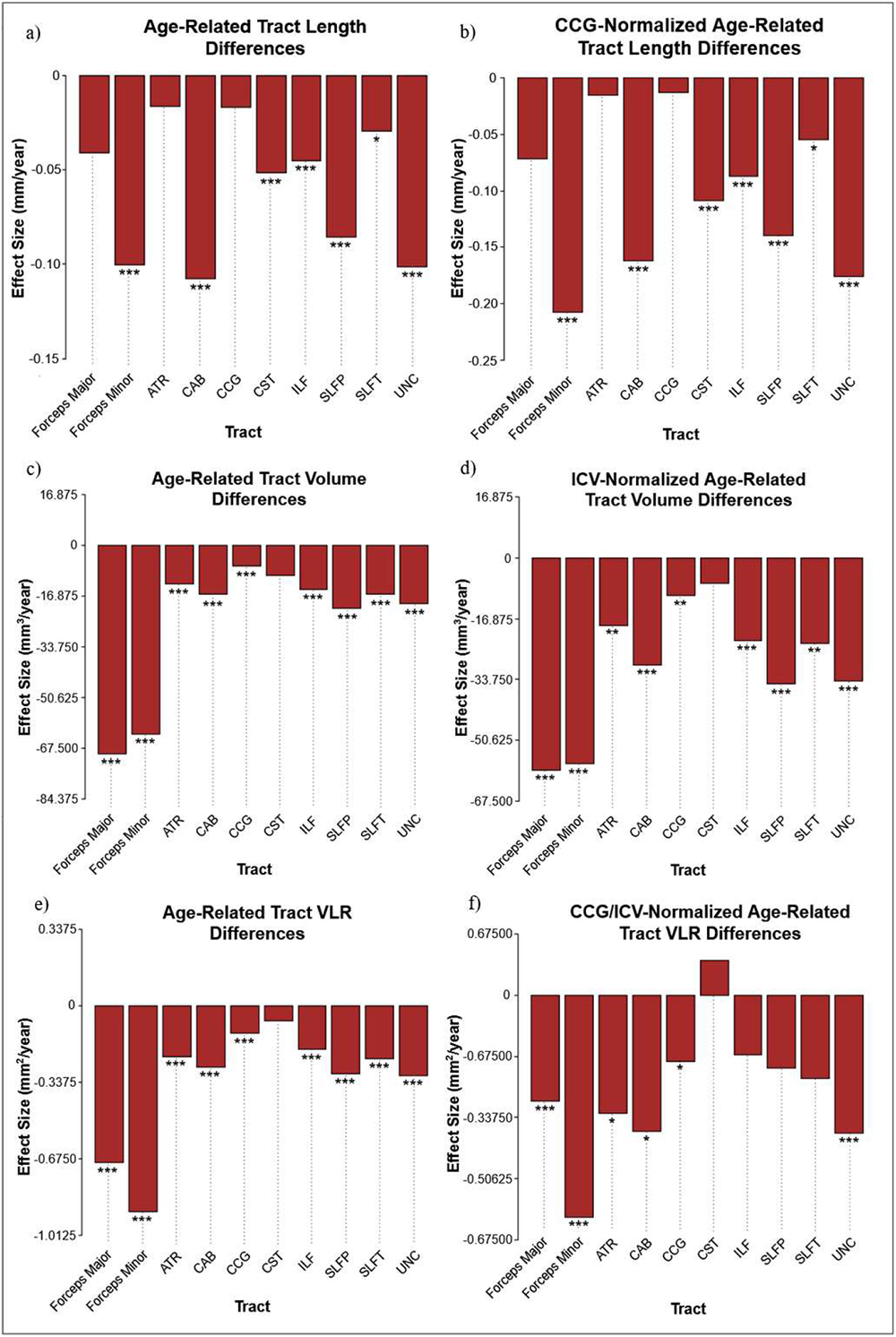
Tract-wise age effects on raw length, volume, and VLR (a, c,e) and Length_norm_, Volume_norm_, and VLR_norm_ (b, d, f). Asterisks denote significant age effects (***: p<.001, **: p<.01, and *: p<.05).

#### Sex Differences

Analysis of raw morphometric values identified significant sex differences in raw tract Length, raw tract Volume, and raw tract VLR for all ten tracts, with male subjects showing higher values in all three metrics across tracts. Female subjects demonstrated higher tract Length_norm_ in the CST, while male subjects demonstrated higher Length_norm_ in the Fmajor, Fminor, ATR, CCG, ILF, SLFP, SLFT, and UNC. Male subjects also showed significantly higher Volume_norm_ and VLR_norm_ in all ten tracts of interest (**Figure 3**).

**Figure 3:**
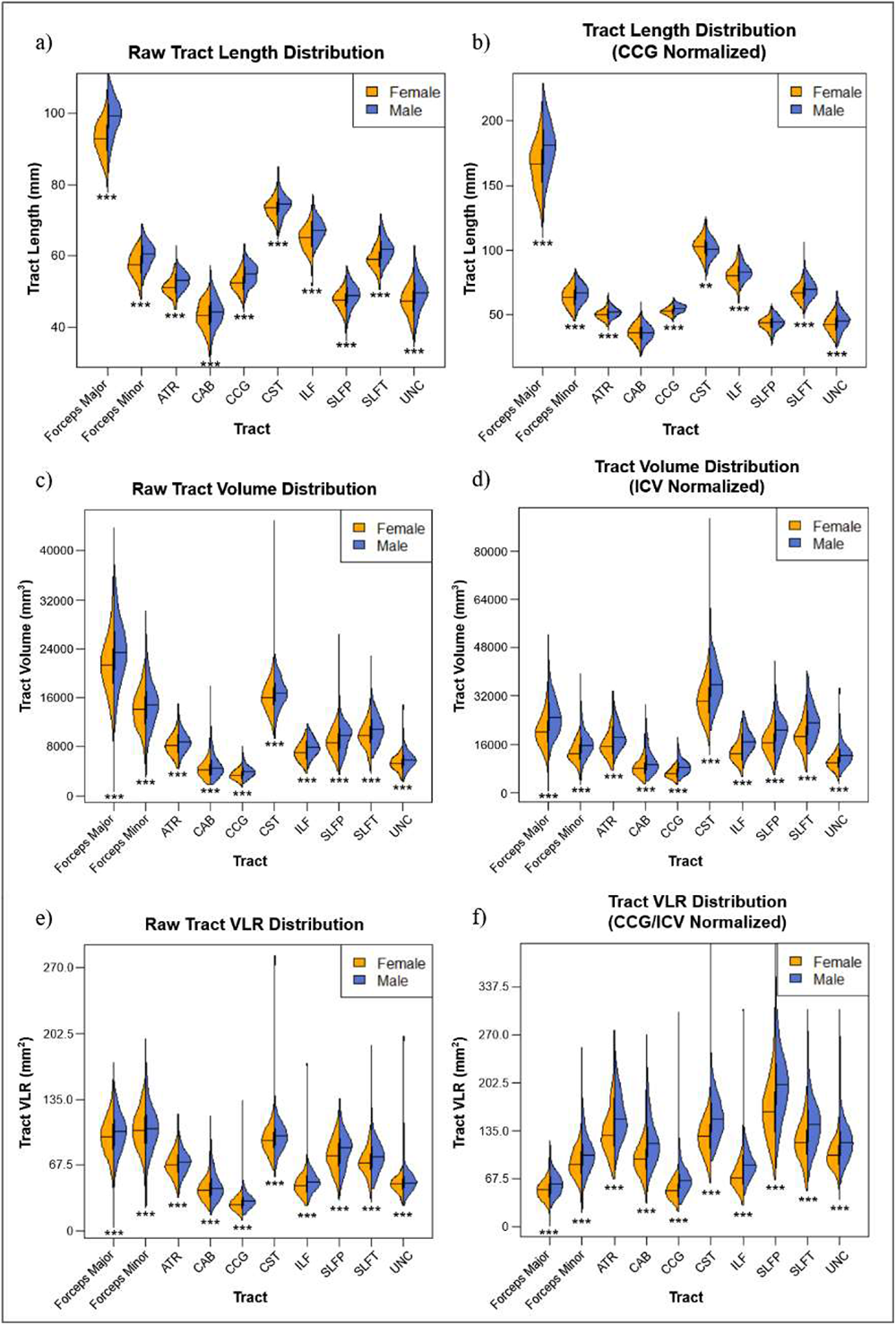
Tract-wise raw Length, Volume, and VLR (a, c,e) and Length_norm_, Volume_norm_, and VLR_norm_ (b, d, f) distributions separated by sex. Asterisks denote significant sex differences (***: p<.001, **: p<.01, and *: p<.05)

### Microstructure

#### MD/FA vs. Age and Sex

All ten tracts of interest showed significant positive relationships between age and MD_perc_. Age was significantly negatively related to FA_perc_ in the Fmajor, Fminor, ATR, CAB, CCG, ILF, and SLFP (**Figure 4**). Female subjects demonstrated significantly higher MD_perc_ in the CST (F(2, 497) = 47.0, p < .001, R^2^_Adjusted_ = .16, effect = 0.000028 mm^2^/s) and in the UNC (F(2, 497) = 71.2, p < .001, R^2^_Adjusted_ = .22, effect = 0.0000076 mm^2^/s). In the ILF, men demonstrated significantly higher MD_perc_ (F(2, 497) = 63.3, p < .001, R^2^_Adjusted_ = .20, effect = 0.0000067 mm^2^/s). In CAB, female subjects demonstrated significantly higher FA_perc_ (F(2, 497) = 14.0, p < .001, R^2^_Adjusted_ = .05, effect = 0.011). In UNC, male subjects demonstrated significantly higher FA_perc_ (F(2, 497) = 3.1, p = .046, R^2^_Adjusted_ = .008, effect = 0.0063). Quadratic, rather than linear regression provided a better fit for age effects on MD_perc_ in all tracts, as well as for age effects on FA_perc_ in Fminor, CCG, and ILF (**Supplemental Figure 2**).

**Figure 4:**
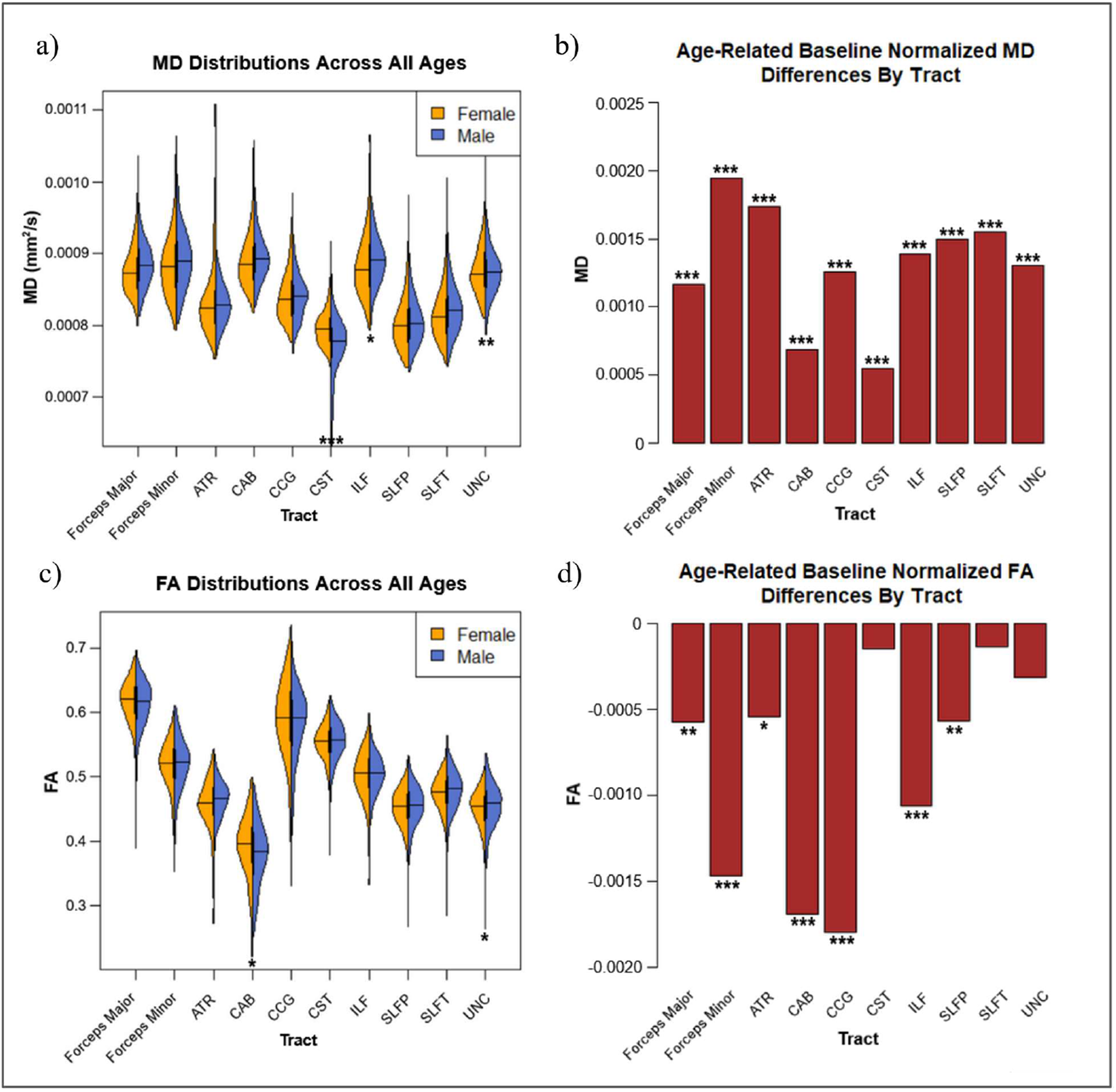
Raw MD and FA distributions separated by sex (a,c), and tract-wise age effects for MD_perc_, and FA_perc_ (b,d). Significant effects by sex and age are denoted with asterisks (***: p<.001, **: p<.01, and *: p<.05)

### Morphometry-Microstructure Relationships

#### Model 1 (morphometry-microstructure relationships)

Mean tract Length_perc_ was significantly negatively associated with MD_perc_ (F(2, 497) = 12.3, p < .001, R^2^_Adjusted_ = .04, slope = -0.21), and significantly positively associated with FA_perc_ (F(2, 497) = 8.1, p < .001, R^2^_Adjusted_ = .03, slope = 0.27). Mean tract VLR_perc_ was significantly negatively associated with both MD_perc_ (F(2, 497) = 11.3, p < .001, R^2^_Adjusted_ = .04, slope = -0.049) and FA_perc_ (F(2, 497) = 8.4, p < .001, R^2^_Adjusted_ = .03, slope = -0.069) (**Supplemental Table 1a**).

Length_perc_ vs. MD_perc_ and FA_perc_: significant negative associations with MD_perc_ were identified in Fminor, CST, SLFP and UNC. Significant positive associations with FA_perc_ were identified in Fminor, ATR, CAB, CST, and ILF, and SLFT.

VLR_perc_ vs. MD_perc_ and FA_perc_: significant negative associations with MD_perc_ were identified in the Fmajor, Fminor, ATR, CCG, SLFP, SLFT, and UNC, and a significant positive association in the CAB. Significant negative associations with FA_perc_ were identified in the Fmajor, ATR, CAB, CST, and UNC (**Supplemental Table 1b**).

#### Model 2 (Sex differences in morphometry-microstructure relationships)

After dividing the sample by sex, a significant positive association was found between Length_perc_ and FA_perc_ in the female sample (F(2, 291) = 13.2, p < .001, R^2^_Adjusted_ = .08, slope = 0.26), while negative associations were identified between VLR_perc_ and FA_perc_ in both female (F(2, 291) = 23.5, p < .001, R^2^_Adjusted_ = .13, slope = -0.13) and male (F(2, 203) = 10.7, p < .001, R^2^_Adjusted_ = .09, slope = -0.11) subjects. No significant associations were identified between morphometry and MD_perc_ after dividing the sample by sex (**Supplemental Table 2a**).

Length_perc_ vs. MD_perc_ and FA_perc_: a significant positive association with MD_perc_ was identified in female subjects in the ATR, while a significant negative association was found in the CST for male subjects. Female subjects demonstrated significant positive relationships between length and FA_perc_ in the ATR, CAB, CST, ILF, and SLFT, in addition to a significant negative association in the UNC, while male subjects demonstrated a significant negative association in the SLFP and a significant positive association in the ILF.

VLR_perc_ vs. MD_perc_ and FA_perc_: both female and male subjects demonstrated significant positive associations between VLR and MD_perc_ in the CAB, with male subjects showing significant negative associations in the Fmajor and SLFT. Both sexes demonstrated significant negative associations between VLR and FA_perc_ in the Fmajor, Fminor, ATR, CAB, CST, and UNC, with Female subjects showing significant negative associations in the ILF. Female subjects showed an additional significant positive association with FA_perc_ in the SLFT (**Supplemental Table 2b**).

### Tract-wise canonical correlations

CCA analysis identified varying patterns of tract-wise microstructural and morphometric loadings between male and female subjects (**Figure 5**). Loadings are listed only for statistically significant canonical correlations, and all loadings are thresholded at 0.5. CCA findings are categorized into “morphometric”, “microstructural” and “morphometic-microstructural”, as shown in **Table 1** along with their likely interpretations. Notably, the most common morphometric pattern is (i), which implies prolonged survival of the largest fibres. Sex-independent common morphometric relationships include opposite loadings in length and volume in the ATR, CAB, and SLFT, and volume loading without significant length loading in the CST. Regarding microstructural findings, both males and females exhibit relationship (iv) in the UNC and pattern (vi) in the ATR, implying degeneration of crossing fibres in these tracts. Both sexes also exhibit pattern (vii) in the SLFP, suggesting negligible crossing-fibre contribution.

**Figure 5:**
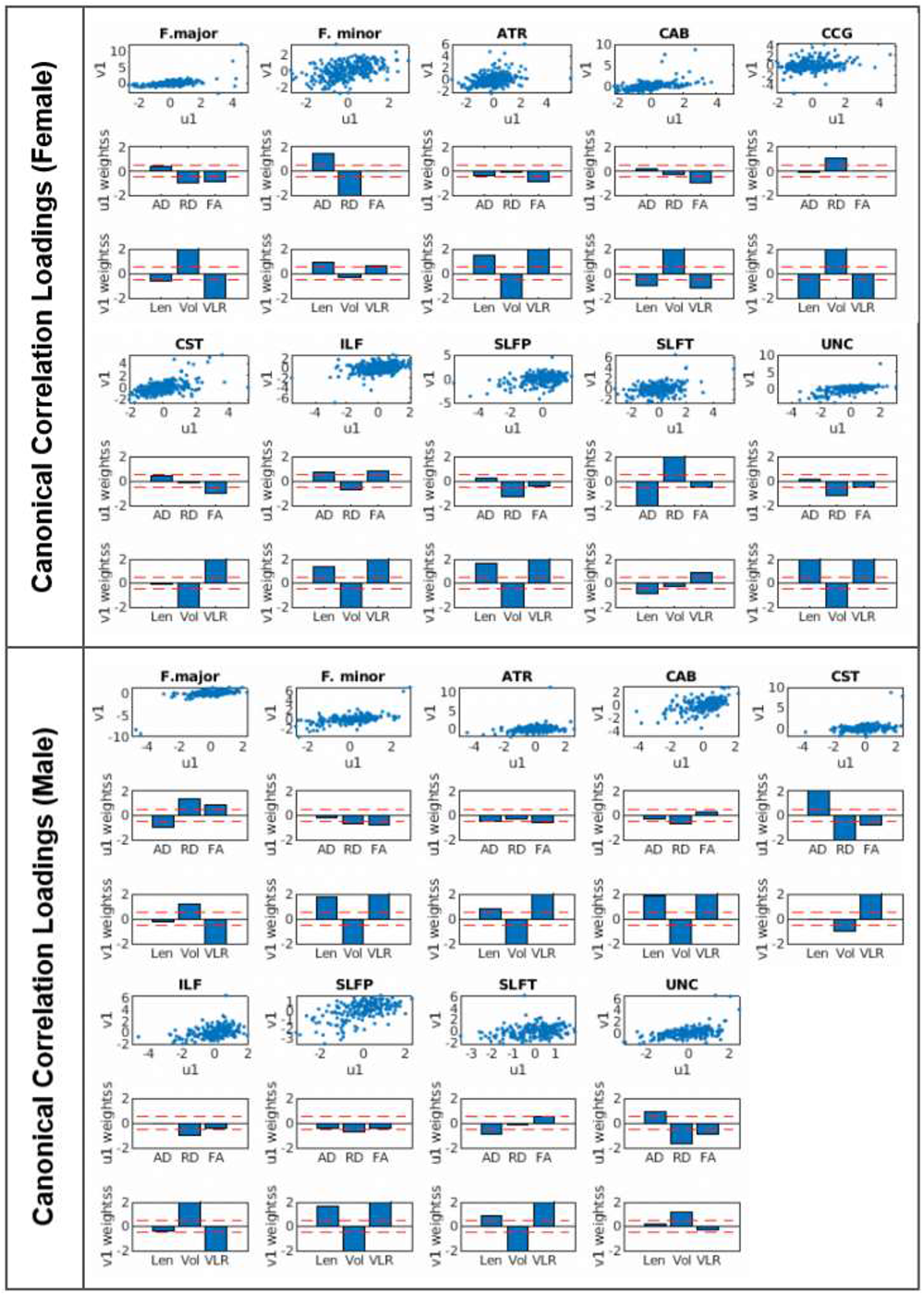
Tract-wise contributions by microstructural (AD_perc_/RD_perc_/FA_perc_) and morphometric (Length_perc_/Volume_perc_/VLR_perc_) measures to categorical perfusion-microstructure relationships separated by sex. Dotted red lines denote the loading threshold (i.e. 0.5).

The morphometric-microstructural patterns displayed no commonality between sexes. Males exhibit pattern (ii) in 3 additional tracts than females (i.e. Fmajor, ILF, UNC), implying less participation of secondary fibre degeneration in the morphometric loss. Females display pattern (vii) in anterior-superior WM (i.e. Fminor, CCG), a sign of negligible cross-fibre contribution, whereas males display pattern (vii) in lateral-inferior tracts (i.e. CAB, ILF). Females exhibit more significant microstructural-morphometric loading patterns than males. Females exhibit pattern (viii) in the Fminor and SLFP, implicating loss of tract length and packing density, consistent with their display of pattern (iii); males do not. Instead, males exhibit pattern (ix) in the Fmajor, also associated with loss of packing density. Males alone exhibit pattern (x) in the UNC, associated with more advanced degeneration.

**Table 1:** Histological interpretations of patterns in the morphometry-microstructure relationships identified by the CCA.

| Category | Pattern | Likely interpretation | Implicated tracts |
| --- | --- | --- | --- |
| **Morphometric** | **(i) Len and vol loading opposite** | The largest fibres surviving longest in spite of tract atrophy, intermediate degeneration | F(ATR, CAB, SLFP, Fmajor, ILF, UNC, CCG)<br>M(ATR, CAB, SLFP, Fminor, SLFT) |
|  | **(ii) Vol loading with negligible len** | Fibers degenerate evenly, regardless of size | F(CST)<br>M(CST, Fmajor, ILF, UNC) |
|  | **(iii) Len loading with negligible vol** | Fibre degeneration driven by loss in length and fibre packing density | F(Fminor, SLFT) |
| **Microstructural** | **(iv) RD and FA loading coincide** | Selective degeneration of crossing fibres - secondary fibres degenerate more | F(UNC, Fmajor)<br>M(UNC, Fminor, CST) |
|  | **(v) RD and FA loadings opposite** | Negligible selective degeneration of crossing fibres | F(SLFT) |
|  | **(vi) FA loading with negligible RD** | Crossing fibre contribution - demyelination with obstruction of axial diffusivity, **or** primary fibre contribution - obstruction of axial diffusivity with minimal demyelination | F(ATR, CAB, CST)<br>M(ATR, SLFT) |
|  | **(vii) RD loading with negligible FA** | Selective degeneration of crossing fibres - secondary fibres degenerate more | F(SLFP, Fminor, CCG)<br>M(SLFP, CAB, ILF) |
| **Morphometric-microstructural** | **(viii) Len and RD loadings opposite** | Loss in length co-exists with rise in extracellular space, implicating loss of packing density | F(Fminor, SLFT) |
|  | **(ix) Vol and RD loadings together** | Volume is higher in the presence of loss of packing density driven by increase in extracellular space | M(Fmajor) |
|  | **(x) Vol and RD loadings opposite** | Loss in volume co-exists with rise in extracellular space, more advanced degeneration | F(Fmajor)<br>M(UNC) |

### Tract-wise cross-correlations

At zero lag, significant correlations between Length_perc_ and FA_perc_ were identified in the Fminor, and SLFP, between Length_perc_ and AD_perc_ in the Fminor, ATR, CAB, CCG, ILF, and SLFP, and between Length_perc_ and RD_perc_ in the Fminor, ATR, CAB, CCG, ILF, and SLFP. A single correlation at negative lag was identified between Length_perc_ and FA_perc_ in the CCG, in which Length_pert_ led FA_perc_ by approximately seven years. No significant tract-wise correlations were identified between Volume_perc_ and our microstructural variables.

Comparing Length_perc_ to Volume_perc_ we found significant correlations at 0 lag in Fminor, ATR, CCG, ILF, and SLFP. All tracts demonstrated significant 0 lag peak correlations between AD_perc_ and RD_perc_. Significant correlations between both AD_perc_ and FA_perc_, as well as AD_perc_ and FA_perc_, were identified at 0 lag in Fminor, CCG, CST, SLFP, and UNC, along with correlations at negative lag in the SLFT, in which the peak CCF saw both AD_perc_ and RD_perc_ leading FA_perc_ by approximately 14 years (**Figure 6**).

**Figure 6:**
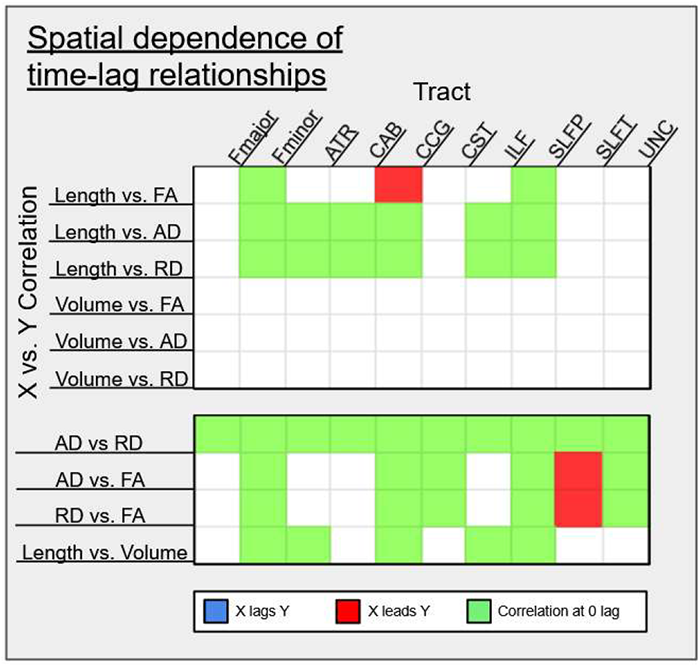
Tract-wise cross-correlation results demonstrating significant (p<.05) peak correlations at positive, negative and zero lag by age category.

## DISCUSSION

Our initial hypotheses included that (1) age-related microstructural differences would be greater in tracts that demonstrate greater age-related morphometrical differences, and (2) tracts that become relatively thinner with age will have relatively high mean tract lengths. The age effects on morphometry and microstructure are summarized in **Figure 7** for comparison. While consistent trends in morphometric and microstructural declines with age were identified in the majority of tracts, this was not the case for the morphometry-microstructure relationships. That is, unlike hypothesized, tract-wise relationships between morphometry and microstructure were demonstrated to be highly variable across tracts as well as between sexes.

**Figure 7:**
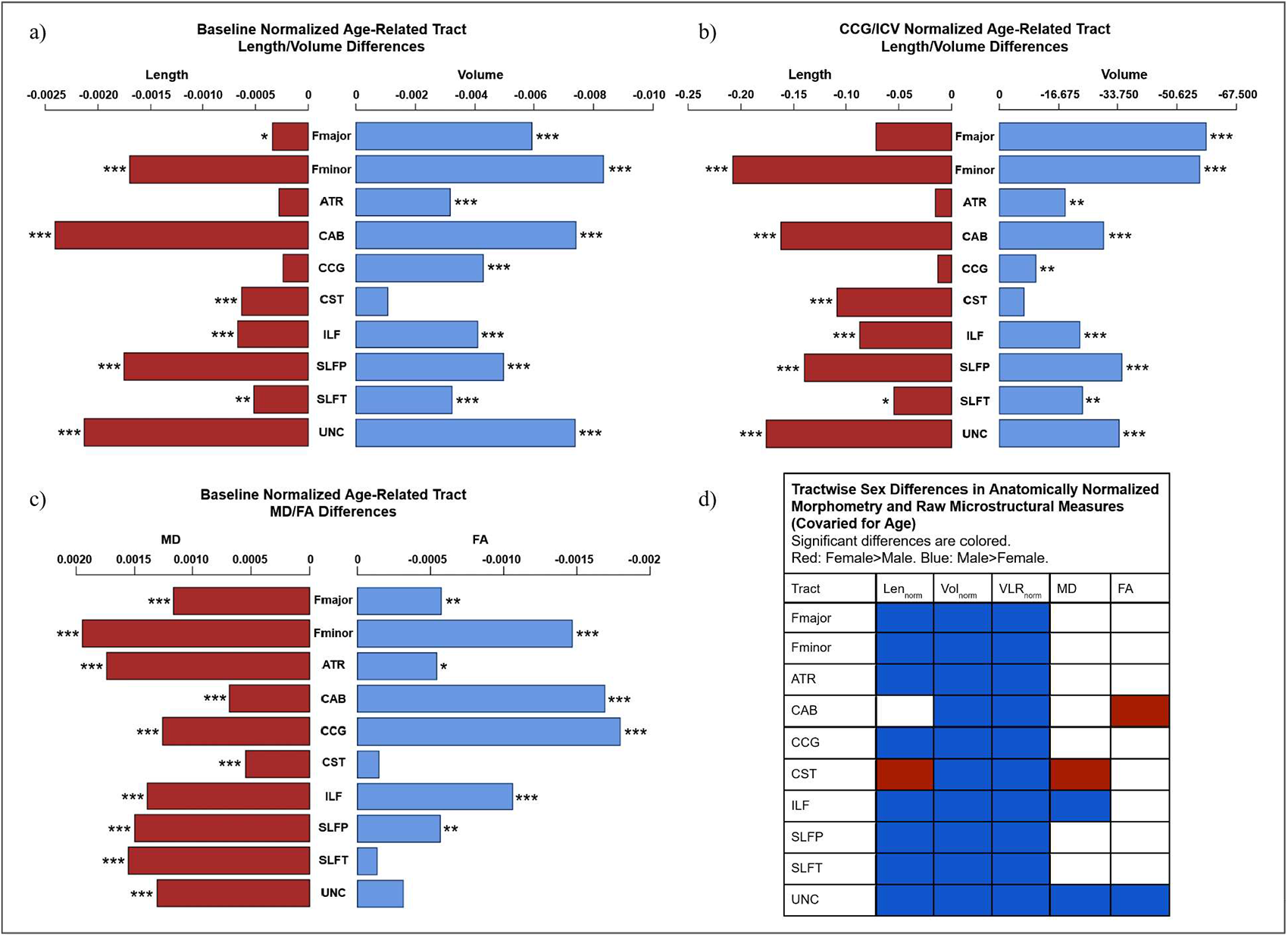
Summary of tract-wise age effects for Length_perc_ and Volume_perc_ (a), Length_norm_ and Volume_norm_ (b), and MD_perc_ and FA_perc_ (c), and sex differences in Length_norm_, Volume_norm_, VLR_norm_, MD, and FA, controlled for age effects (d). Asterisks denote significant age effects (***: p<.001, **: p<.01, and *: p<.05).

### Morphometry and age

In line with existing research on white matter atrophy (de Groot et al., 2015; Sala et al., 2012; Tang et al., 1997), we observed broad age-related decreases in all three raw morphometric measures. At the tract-wise level, decreasing raw tract volume in aging was found in all tested tracts with the exception of the CST. Conversely to the CST, the major forceps and minor forceps exhibited notably greater age-related raw volume differences than other WM regions. These findings corroborate recent work noting broad morphometric declines across the brain, with commissural tracts, including both the forceps major and minor, as pathways with relatively high susceptibility to age-related change, and the CST relatively preserved (Schilling et al., 2022). All tracts except three (major forceps, ATR, and CCG) also shortened in aging (**Figure 2**). The CCG, due to the observed age stability in its length, recommended itself as the normalization target in subsequent tract length analyses. When length and volume were both normalized (volume by ICV and length by CCG length), the length_perc_ and volume_perc_ decreases in WM overall still outpaced those in the reference structure.

Taken together, our findings demonstrate that length_perc_ and volume_perc_ do not decline proportionately in aging. Tracts such as the CST do in fact shorten with age without significant proportionate losses in volume. In contrast, tracts such as the CCG would be expected to demonstrate the opposite pattern, thinning with age as it loses volume_perc_ with relatively little loss in tract length_perc_. Another interpretation is that tract-wise morphometric variability represents a cross-sectional observation of stages in an ongoing, but shared trajectory of decline. This would suggest that the pattern of morphometric decline is common across the brain, but takes effect at different ages or rates between tracts due to regionally specific protective or exacerbating factors.

### Microstructure and age

Relationships between age and WM integrity across tracts note the expected trends of increasing MD_perc_ and decreasing FA_perc_ with advancing age. This echoes general findings in the literature regarding microstructural WM decline (Benitez et al., 2018; Sala et al., 2012). As in morphometric age associations, tracts varied considerably in their trajectory of microstructural changes. While all tracts demonstrated increases in MD, three tracts, namely the CST, SLFT, and UNC, exhibited no significant effects of age on FA_perc_. A possible source of this variability between microstructural metrics is the contribution of crossing fibers to diffusion measures within a tract, such as regional increases in FA due to degeneration of crossing fibers, despite relatively maintained MD. Moreover, increasing AD can potentially be due to the deterioration of crossing fibers resulting in RD increase of the secondary (perpendicular) fibre direction, which translates to AD increases in the principal fibre direction (Chad et al., 2018; Han et al., 2023). This pattern between tracts also continues the trend observed in the morphometric data, in which the CST shows considerable robustness to aging declines. As both the SLFT and UNC are relatively posterior tracts, these findings may be interpretable as support for retrogenesis theories on brain aging, however, significant MD_perc_ increases with age were nonetheless identified in all three tracts. Moreover, the UNC showed strong age effects on morphometry in both length_perc_ and volume_perc_. Prior research has similarly noted considerable heterogeneity in macrostructural age changes across the WM in spite of more homogenous rates of microstructural deterioration (Schilling et al., 2022), demonstrating the need to examine relationships between morphometry and microstructure to further clarify the basis of tract-wise differences.

### Understanding types of WM aging through multimodal integration

Variation in both relative tract volume_perc_ and length_perc_ are assumed to be the result of age-related culling of axonal fibers, as noted in histological examination of the WM across age (Choy et al., 2020; Marner et al., 2003). Our results demonstrate that these values do not, however, decline together at the same rates across the brain. Variation in WM volume_perc_ across age is present regardless of changes in tract length_perc_, meaning that some tracts show relatively preserved volume_perc_ despite notable losses in tract length_perc_. This is theorized to be the result of variation in extracellular space at different stages of WM decline. Increased size of surviving fibres, as well as thickening of myelin in aging axons, results in notably larger mean diameter in the aging tract (Tang et al., 1997). The geometry of packing larger fibres produces an increase in extracellular space between the fibres and relatively lower impact on tract volume. This is in contrast to even degeneration of both large and small fibers, which could allow a more densely packed bundle and produces relatively greater losses in tract volume (Marner et al., 2003; Tang et al., 1997).

### Sex differences in morphometry-microstructural variations in aging

Sex differences in the morphometric and microstructural parameters are summarized in **Figure 7d**. For the most part, males have higher tract volume, length and VLR, with the exception of the CST, regardless of anatomically normalization. Conversely, there are far fewer sex differences in microstructure. Results from Models 1 and 2 indicate that the micro-macrostructural associations are influenced by age and sex in a tract-specific manner, which was further clarified by the CCA results.

The fusion of morphometric and microstructural observations in CCA allows us to examine regional differences in WM aging in view of expected sex differences. For each combination, we have provided a likely interpretation based on known literature, as summarized in **Supplemental Table 3**. The predominance of fibers demonstrating opposing loading in length_perc_ and volume_perc_ support our study’s initial hypothesis that the culling of vulnerable axon fibers would result in proportionally longer surviving fibers (pattern (i) in the CCA results). Conversely, preferential culling of shorter fibers in relatively posterior tracts (Fminor and SLFT), which lose a greater degree of fiber density rather than fiber volume_perc_ may represent an intermediate step of decline in which the preferential die-off of shorter fibers has begun, but the tract volume_perc_ has not yet compressed. (**Table 1**)

A number of commonalities between sexes were uncovered by the CCA. The CST was notable as the one tract in which relatively preserved length_perc_ was observed in both sexes (pattern (ii)). Other notable commonalities between sexes were the UNC, ATR, and SLFP. In the UNC, both male and female subjects demonstrated relatively lower RD_perc_ in the presence of lower FA_perc_, suggesting contributions by crossing fibers affecting local measures of RD_perc_. A similar argument can be made for commonalities in the ATR between sexes, in which degradation of crossing fibers obstructs diffusivity along the fibers of the tract in question. This could result in FA_perc_ changes despite negligible changes in RD_perc_. Conversely, both male and female subjects displayed contributions by RD_perc_ to microstructural declines in the absence of significant FA loading. In theory, this could indicate that the majority of change in these regions is due to degradation of the tract itself, rather than contributions from crossing fibers.

In addition to the noted similarities, our results suggest sex differences in degree of protection or susceptibility to axonal culling, with microstructural declines in the Fmajor, Fminor, CAB, CST, ILF, and SLFT differing between sexes. As pattern (ii) was more common in males, extending as well to the Fmajor, ILF, and UNC, males may demonstrate less aging-related culling in these regions. Moreover, in male subjects, selective degeneration at crossing fibers appears more prevalent in the Fminor and CST, whereas in females, this is more prevalent in the Fmajor. This may be interpretable as a greater degree of degradation in posterior regions in male subjects, compared to greater prefrontal declines in female subjects. In the SLFT, only male subjects demonstrate patterns of RD_perc_ and FA_perc_ that suggest major contribution by selective degeneration of crossing fibres. Female subjects also demonstrated patterns of RD_perc_ and FA_perc_ that suggest low contribution by selective degeneration. Negligible degeneration in crossing fibers was noted in the female Fminor and CCG, along with the male CAB and ILF, suggesting a fronto-parietal vs temporo-parietal difference between sexes, with these regions showing more advanced declines in female and male subjects, respectively.

As microstructural variations do not distinguish between fibre sizes, the addition of morphometric information is valuable in linking our observations to known histological evidence of the degenerative process. Across tracts, male subjects demonstrate a higher rate of fiber degeneration irrespective of size, notably in the Fmajor, CST, ILF, and UNC. This is in contrast to the preferential degeneration of smaller fibers found across tracts in the female subjects, with the sole exception of the CST. The CST has both shown a higher mean fiber diameter (S. Y. Huang et al., 2020), and been observed to survive relatively intact across the lifespan (de Groot et al., 2016; Yeatman et al., 2014), suggesting that even degeneration of both smaller and larger fibers appears to precede the preferential degeneration by size seen in more advanced stages of decline that results a higher mean tract length_perc_ (Choy et al., 2020; Marner et al., 2003; Tang et al., 1997). Differences in signatures of decline could then reflect differences in range of fiber length, as length_perc_ changes in tracts with relatively uniform fiber length would be more difficult to detect. Determining the presence of sex differences in fiber composition will require further histological examination.

A driving source of this sex discrepancy may also be the hormonal effects of menopause in female subjects, as the impact of menopause on the brain have been demonstrated to include structural, connectivity, and metabolic effects (Mosconi et al., 2021). Our results likely capture structural effects of peri-menopause and post-menopause on the aging WM.

### Temporal relationships in morphometry-microstructural variations in aging

Results from the CCF analysis demonstrated that declines in morphometry and microstructure were largely contemporaneous, with the only notable exception to this pattern being the CCG, in which tract length_perc_ changes appear to lead changes in FA_perc_. As we previously reported, FA is confounded by MD variations, and is thus a sensitive but not specific marker of WM decline. However, this finding may also suggest tract length as a promising target for future studies. As the majority of variations are contemporaneous with one another across categories, this finding could be taken as a representation of the shared mechanisms of decline between the two sets of measures. While this runs counter to the intuitive assumption that microstructural changes should occur prior to morphometric declines, the differences in measurement sensitivities of microstructural and morphometric approaches may complicate the interpretation.

### Limitations

While the intent of this study was to lay the initial groundwork for multimodal integration for understanding white matter aging, the choice to widely applied metrics such as MD and FA introduces a number of limitations that may be resolved in more advanced diffusion modeling. Moreover, this study chose to focus on only the ten major WM tracts. While this provides lower spatial WM coverage than more recent segmentations (Maffei et al., 2021), the choice was made to both minimize the degree of overlap/partial-voluming between tract segmentations, as well as to aid in comparisons with previous research utilizing TRACULA. Additionally, as the CCF depends on nonlinearity in the age-associations of its inputs, the heterogeneity in the presence of significant quadratic versus linear age effects observed across measures restricted the tracts that can be submitted to cross-measure tract-wise comparisons. Finally, while our methods are novel, the cross-sectional nature of the HCP-A dataset merits future validations of our findings using a longitudinal dataset.

## Conclusions

Both morphometry and microstructure analyzed in this study depict WM age-related differences as highly tract-specific, sex specific, and varied across the aging brain. Broad whole-WM effects supported our expectations for normalized volume and length decreases with age, along with anticipated microstructural declines. Examining VLR revealed additional patterns in age-related WM change consistent with our predictions. To our knowledge, this is the first study to incorporate in-vivo WM tract length differences in healthy aging.

Additionally, these findings demonstrate the predominance of crossing WM fibers in the measurement of WM integrity using dMRI. TRACULA provides a valuable tool to map the morphometric changes in our tracts of interest, but the microstructural values measured within those regions are nonetheless influenced by the fibers of other tracts and radiations not included in this model. These contributions must be taken into account when examining tract-based microstructural changes. Our findings suggest that predominantly prefrontal and inferior-frontal regions demonstrate a greater degree of degeneration in crossing WM fibers, notable in our findings in the Fminor, ILF, and UNC, suggesting these regions are broadly more susceptible to decline.

Complicating the task of modeling age-related WM changes is the apparent presence of additional factors in determining which tracts exhibit specific patterns of decline both across and between sexes. These trends cannot be fully accounted for by observed micro-/macrostructural interactions. Additional microstructural and histological variables will need to be incorporated into future models. An area of considerable interest to this study is the exploration of WM perfusion. Imaging of regional perfusion in the WM is a relatively novel capability with observed relationships to WM integrity (Chen et al., 2013; Kim et al., 2020). Regional differences in WM perfusion may represent a missing piece in our understanding of age-related WM declines as a whole.

## Acknowledgments

We are grateful for the financial support from the Canadian Institutes of Health Research (CIHR) (#PJT169688) and the Canada Research Chairs (CRC) program (JJC).

## Supplementary Materials

**Supplementary Table 1a:** Model 1 – Whole-brain relationships between morphometry (length and VLR) and microstructure (covaried for sex). All models are multivariate linear regressions based on baseline-normalized measures (x_perc_) with overall F-statistics, *p*-values, and effect sizes (*R^2^_adjusted_*) listed. Slopes are noted below each regression equation and significant associations bolded with asterisks indicating significance. (*:p<.05, **:p<.01, ***:p<.001).

|  | Baseline Normalized Length |  | Baseline Normalized VLR |  |
| --- | --- | --- | --- | --- |
| Tract | FA | MD | FA | MD |
| All | $F(2, 497) = 8.1, p < .001,$<br>$R^2_{Adjusted} = .03$ | $F(2, 497) = 12.3, p < .001,$<br>$R^2_{Adjusted} = .04$ | $F(2, 497) = 8.4, p < .001,$<br>$R^2_{Adjusted} = .03$ | $F(2, 497) = 11.3, p < .001,$<br>$R^2_{Adjusted} = .04$ |
|  | **0.2739\*\*\*** | **-0.205382\*\*\*** | **-0.069165\*\*\*** | **-0.048962\*\*\*** |

**Supplementary Table 1b:**
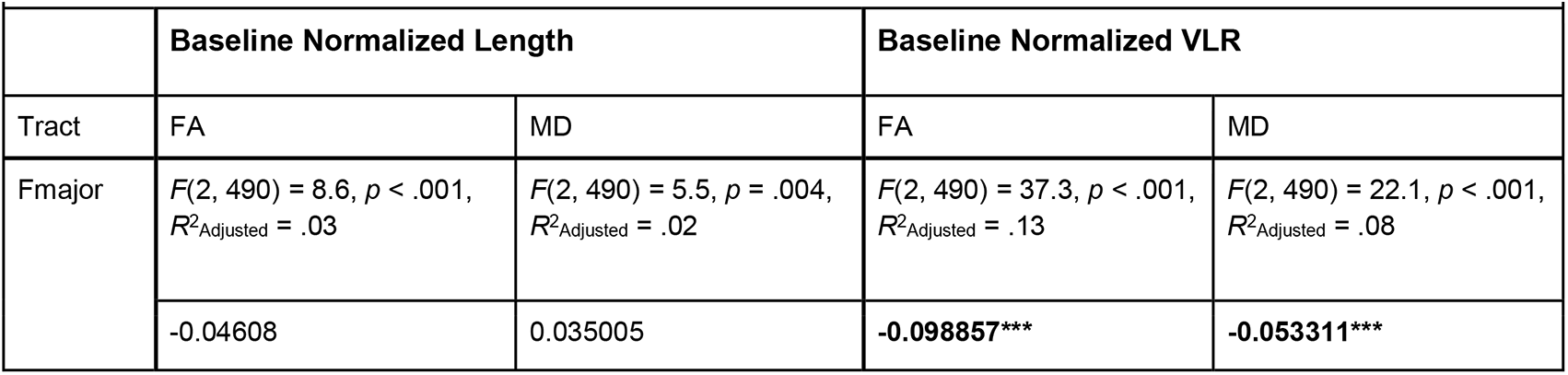
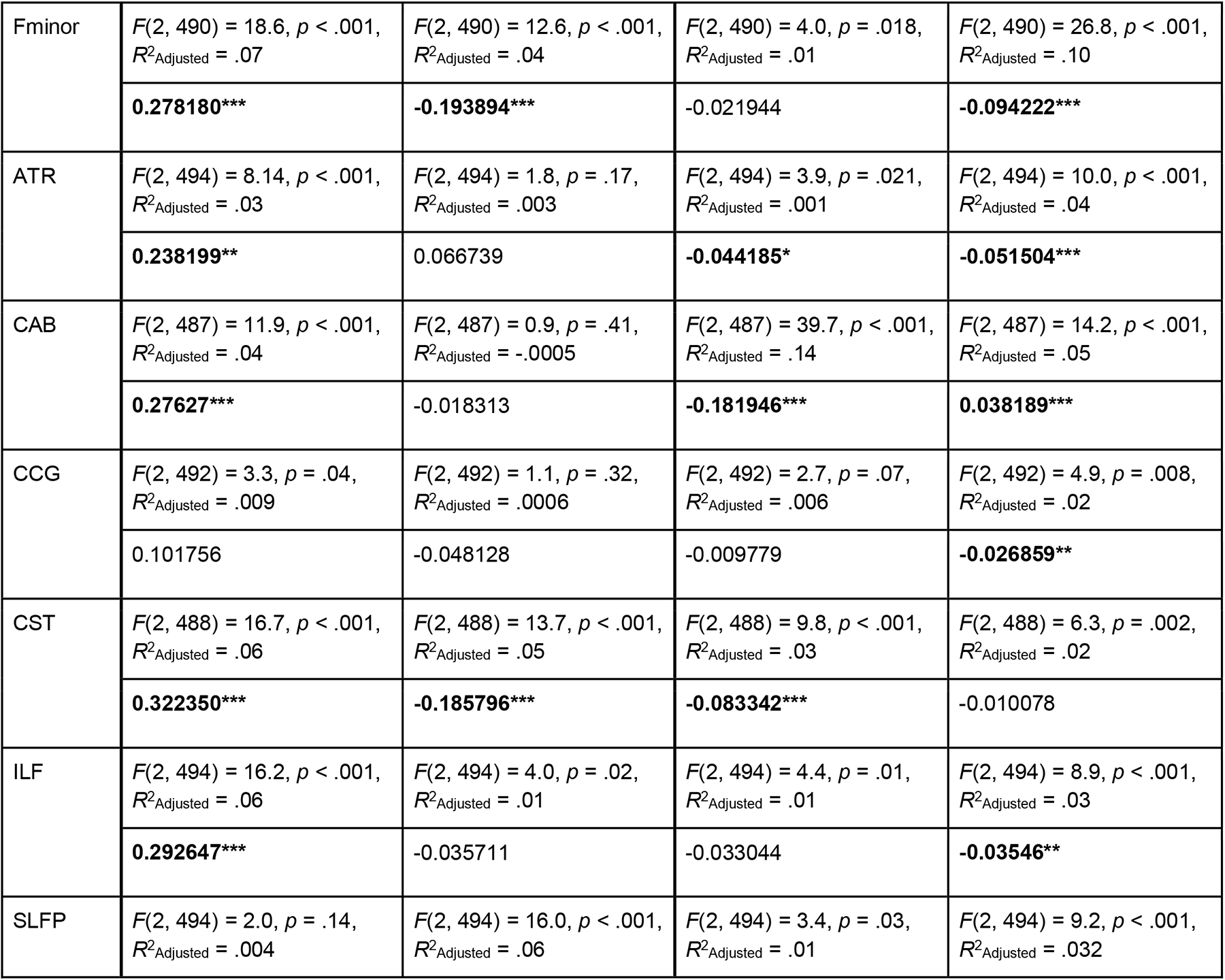
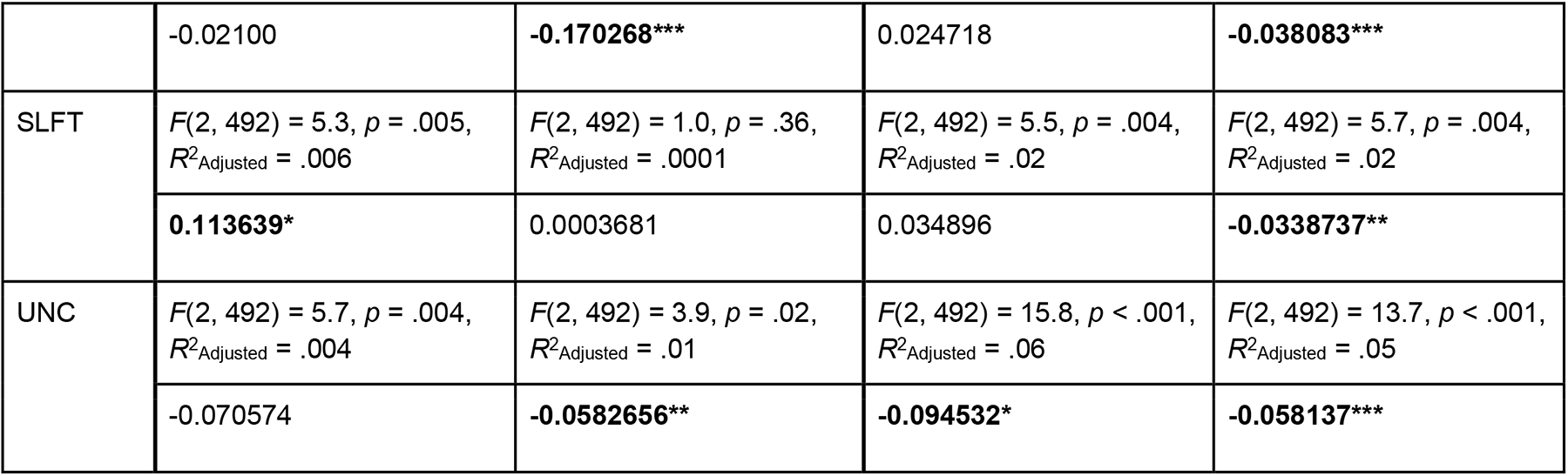
Model 1 – Tractwise relationships between morphometry (length and VLR) and microstructure (covaried for sex). All models are multivariate linear regressions based on baseline-normalized measures (x_perc_) with overall F-statistics, *p*-values, and effect sizes (*R^2^_adjusted_*) listed. Slopes are noted below each regression equation and significant associations bolded with asterisks indicating significance. (*:p<.05, **:p<.01, ***:p<.001).

**Supplementary Table 2a:** Model 2 – Tractwise sex differences in the relationships between morphometry (Length and VLR) and microstructure (covaried for age). All models are multivariate linear regressions based on baseline-normalized measures (x_perc_) with overall F-statistics, *p*-values, and effect sizes (*R^2^_adjusted_*) listed. Slopes are noted below each regression equation and significant associations bolded with asterisks indicating significance. (*:p<.05, **:p<.01, ***:p<.001).

|  | Baseline Normalized Length |  |  |  | Baseline Normalized VLR |  |  |  |
| --- | --- | --- | --- | --- | --- | --- | --- | --- |
|  | FA |  | MD |  | FA |  | MD |  |
| Tract | Female | Male | Female | Male | Female | Male | Female | Male |
| All | $F(2, 291) = 13.2, p < .001, R^2_{Adjusted} = .08$ | $F(2, 203) = 1.7, p = .19, R^2_{Adjusted} = .006$ | $F(2, 291) = 55.0, p < .001, R^2_{Adjusted} = .27$ | $F(2, 203) = 23.3, p < .001, R^2_{Adjusted} = .18$ | $F(2, 291) = 23.5, p < .001, R^2_{Adjusted} = .13$ | $F(2, 203) = 10.7, p < .001, R^2_{Adjusted} = .09$ | $F(2, 291) = 55.2, p < .001, R^2_{Adjusted} = .27$ | $F(2, 203) = 23.3, p < .001, R^2_{Adjusted} = .18$ |
|  | **0.2642212\*\*** | 0.0860415 | 0.0173169 | -0.0825040 | **-0.1276850\*\*\*** | **-0.1078291\*\*\*** | 0.0091113 | -0.0128556 |

**Supplementary Table 2b:**
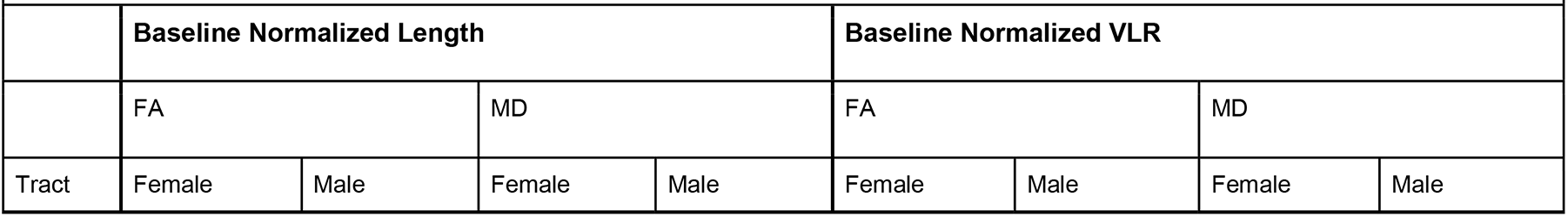
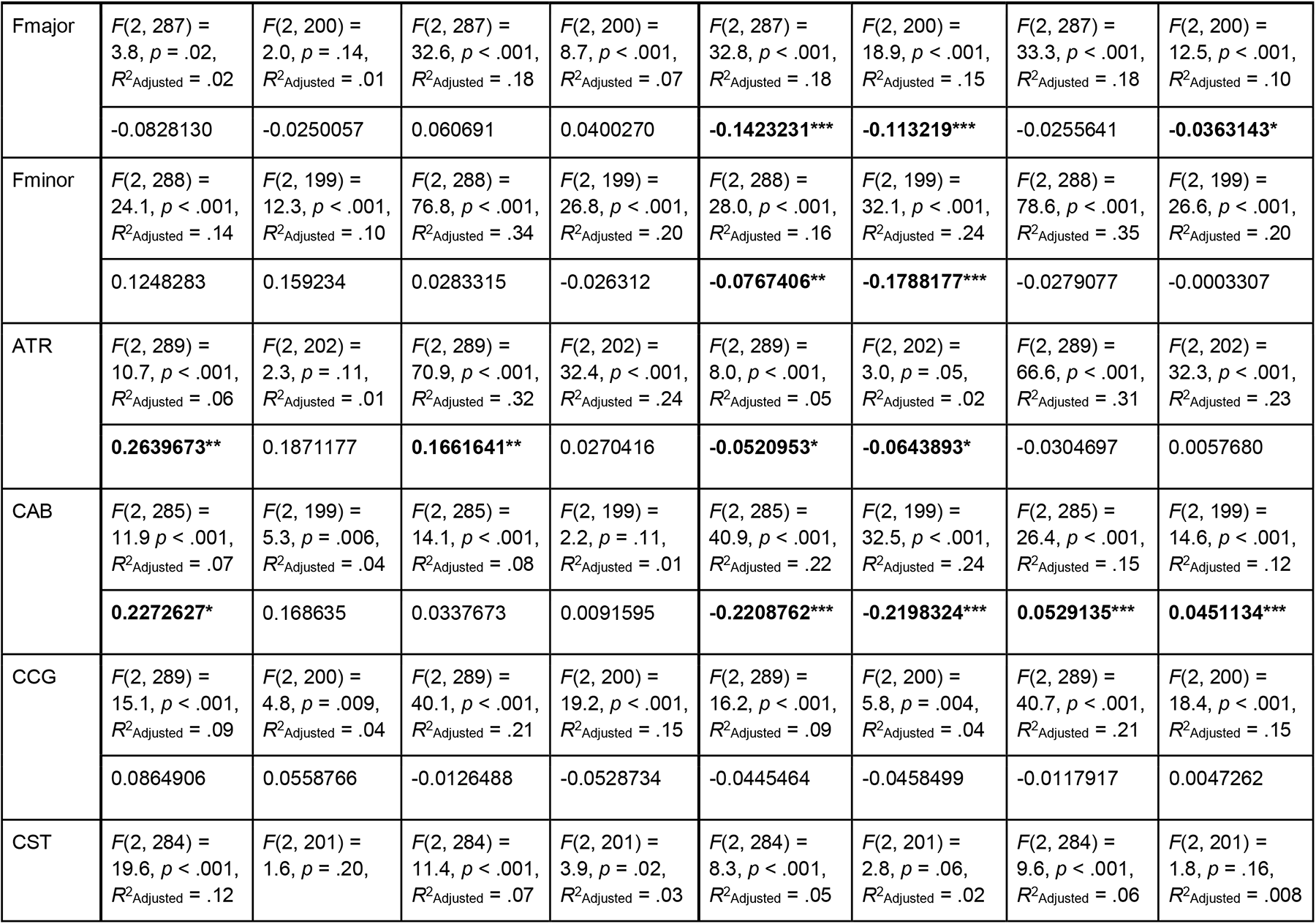
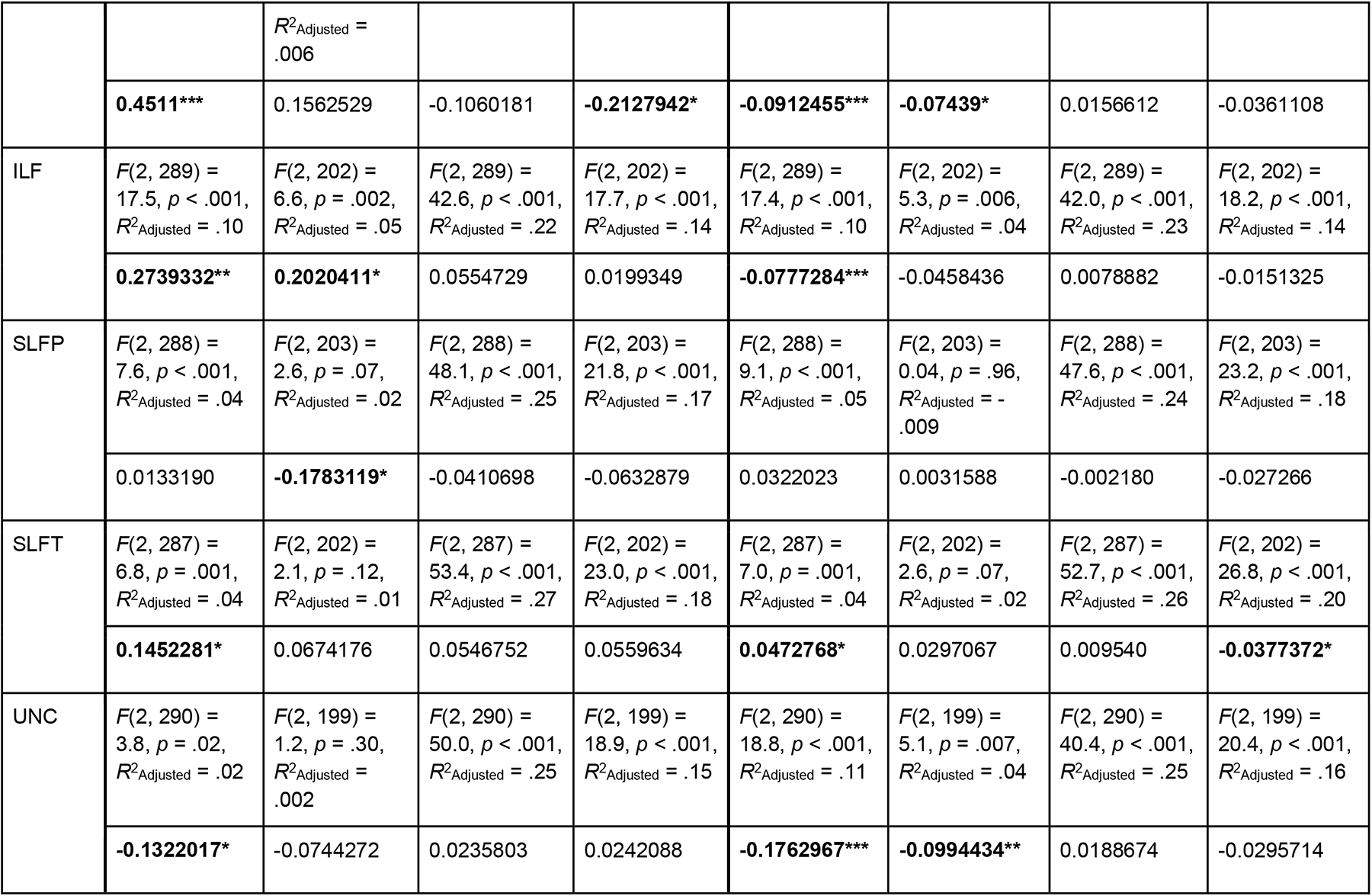
Model 2 – Tractwise sex differences in relationships between morphometry (length and VLR) and microstructure (covaried for age). All models are multivariate linear regressions based on baseline-normalized measures (x_perc_) with overall F-statistics, *p*-values, and effect sizes *R^2^_adjusted_* listed. Slopes are noted below each regression equation and significant associations bolded with asterisks indicating significance. (*:p<.05, **:p<.01, ***:p<.001).

**Supplementary Table 3:** Proposed mechanisms of degeneration identified in each tract of interest based on results from CCA analysis. (Taha et al., 2022; Uddin et al., 2019)

|  | **Female** | **Male** |
| --- | --- | --- |
| **Fmajor** | Prolonged survival of largest fibres with maintained packing density. Selective degeneration of crossing fibres. | Even degeneration by fiber size with maintained packing density, with rise in extracellular space |
| **Fminor** | Prolonged survival of largest fibres with loss of packing density. Selective degeneration of crossing fibres. | Prolonged survival of largest fibres with maintained fiber density. Selective degeneration of crossing fibres. |
| **ATR** | Prolonged survival of largest fibres with maintained packing density. Crossing fiber demyelination with AD obstruction. | Prolonged survival of largest fibres with maintained packing density. Crossing fiber demyelination with AD obstruction. |
| **CAB** | Prolonged survival of largest fibres with maintained packing density. Crossing fiber demyelination with AD obstruction. | Prolonged survival of largest fibres with maintained packing density. Selective degeneration of crossing fibres. |
| **CCG** | Prolonged survival of largest fibres with maintained packing density. Selective degeneration of crossing fibres. | No association between morphometry and microstructure. |
| **CST** | Even degeneration by fiber size with maintained packing density. Crossing fiber demyelination with AD obstruction. | Even degeneration by fiber size with maintained packing density. Selective degeneration of crossing fibres. |
| **ILF** | Prolonged survival of largest fibres with maintained packing density. Negligible crossing fiber degeneration. | Even degeneration by fiber size with maintained packing density. Selective degeneration of crossing fibres. |
| **SLFP** | Prolonged survival of largest fibres with maintained packing density. Selective degeneration of crossing fibres. | Prolonged survival of largest fibres with maintained packing density. Selective degeneration of crossing fibres. |
| **SLFT** | Prolonged survival of largest fibres with loss of packing density. Negligible crossing fiber degeneration. | Prolonged survival of largest fibres with maintained packing density. Crossing fiber demyelination with AD obstruction. |
| **UNC** | Prolonged survival of largest fibres with loss of packing density. Selective degeneration of crossing fibres. | Even degeneration by fiber size with maintained packing density. Selective degeneration of crossing fibres. |

**Supplementary Figure 1:**
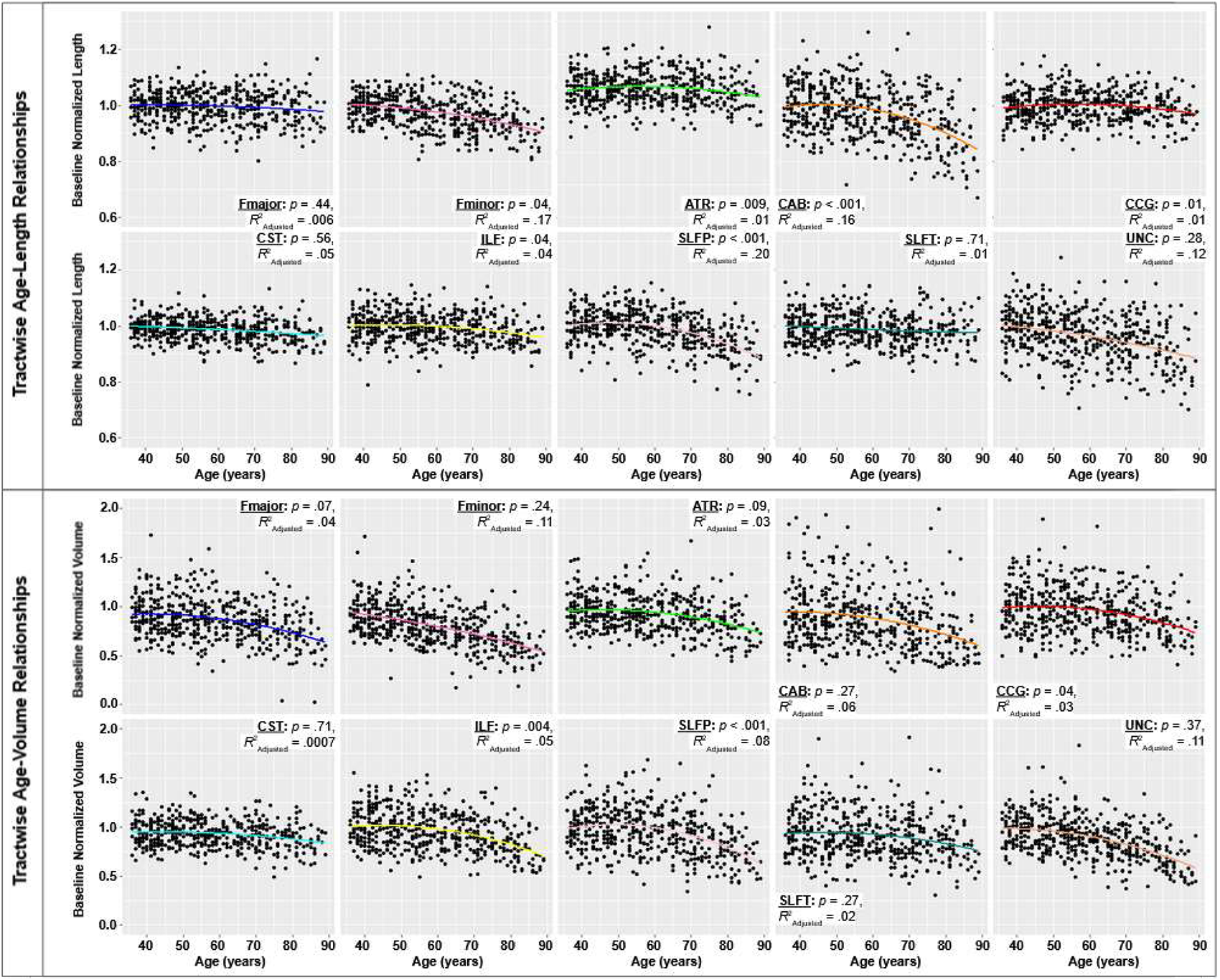
Tractwise quadratic regression of age effects on Length_perc_ and Volume_perc_. P-values and effect sizes from quadratic regression are noted for each tract.

**Supplementary Figure 2:**
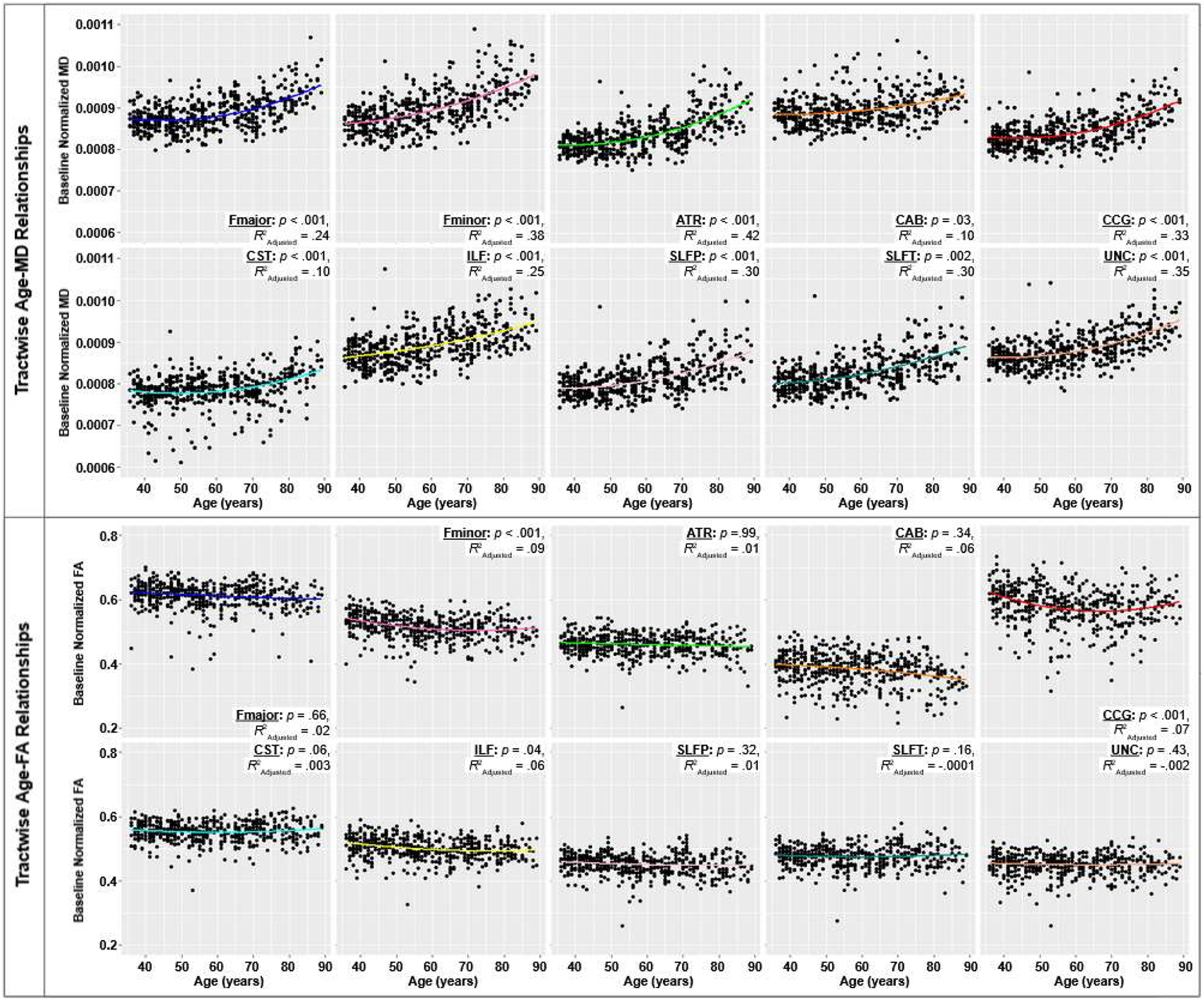
Tractwise regression of Length_perc_ and Volume_perc_ against age with each dot representing one subject. The p-values and effect sizes (R^2^_adjusted_) from the quadratic fit are noted for each tract.

## REFERENCES

Armstrong, C. L., Traipe, E., Hunter, J. V., Haselgrove, J. C., Ledakis, G. E., Tallent, E. M., Shera, D., & van Buchem, M. A. (2004). Age-related, regional, hemispheric, and medial-lateral differences in myelin integrity in vivo in the normal adult brain. AJNR. American Journal of Neuroradiology, 25(6), 977–984.

Armstrong, N. M., An, Y., Shin, J. J., Williams, O. A., Doshi, J., Erus, G., Davatzikos, C., Ferrucci, L., Beason-Held, L. L., & Resnick, S. M. (2020). Associations between cognitive and brain volume changes in cognitively normal older adults. NeuroImage, 223, 117289.

Avellino, A. M., Hart, D., Dailey, A. T., MacKinnon, M., Ellegala, D., & Kliot, M. (1995). Differential macrophage responses in the peripheral and central nervous system during wallerian degeneration of axons. Experimental Neurology, 136(2), 183–198.

Azevedo, C. J., Cen, S. Y., Jaberzadeh, A., Zheng, L., Hauser, S. L., & Pelletier, D. (2019). Contribution of normal aging to brain atrophy in MS. Neurology(R) Neuroimmunology & Neuroinflammation, 6(6). 10.1212/NXI.0000000000000616

Bajada, C. J., Schreiber, J., & Caspers, S. (2019). Fiber length profiling: A novel approach to structural brain organization. NeuroImage, 186, 164–173.

Baker, L. M., Laidlaw, D. H., Cabeen, R., Akbudak, E., Conturo, T. E., Correia, S., Tate, D. F., Heaps-Woodruff, J. M., Brier, M. R., Bolzenius, J., Salminen, L. E., Lane, E. M., McMichael, A. R., & Paul, R. H. (2017). Cognitive reserve moderates the relationship between neuropsychological performance and white matter fiber bundle length in healthy older adults. Brain Imaging and Behavior, 11(3), 632–639.

Baker, L. M., Laidlaw, D. H., Conturo, T. E., Hogan, J., Zhao, Y., Luo, X., Correia, S., Cabeen, R., Lane, E. M., Heaps, J. M., Bolzenius, J., Salminen, L. E., Akbudak, E., McMichael, A. R., Usher, C., Behrman, A., & Paul, R. H. (2014). White matter changes with age utilizing quantitative diffusion MRI. Neurology, 83(3), 247–252.

Bartzokis, G. (2004). Age-related myelin breakdown: a developmental model of cognitive decline and Alzheimer’s disease. Neurobiology of Aging, 25(1), 5–18; author reply 49–62.

Bastin, M. E., Muñoz Maniega, S., Ferguson, K. J., Brown, L. J., Wardlaw, J. M., MacLullich, A. M. J., & Clayden, J. D. (2010). Quantifying the effects of normal ageing on white matter structure using unsupervised tract shape modelling. NeuroImage, 51(1), 1–10.

Benitez, A., Jensen, J. H., Falangola, M. F., Nietert, P. J., & Helpern, J. A. (2018). Modeling white matter tract integrity in aging with diffusional kurtosis imaging. Neurobiology of Aging, 70, 265–275.

Bookheimer, S. Y., Salat, D. H., Terpstra, M., Ances, B. M., Barch, D. M., Buckner, R. L., Burgess, G. C., Curtiss, S. W., Diaz-Santos, M., Elam, J. S., Fischl, B., Greve, D. N., Hagy, H. A., Harms, M.P., Hatch, O. M., Hedden, T., Hodge, C., Japardi, K. C., Kuhn, T. P., … Yacoub, E. (2019). The Lifespan Human Connectome Project in Aging: An overview. NeuroImage, 185, 335–348.

Brickman, A. M., Meier, I. B., Korgaonkar, M. S., Provenzano, F. A., Grieve, S. M., Siedlecki, K. L., Wasserman, B. T., Williams, L. M., & Zimmerman, M. E. (2012). Testing the white matter retrogenesis hypothesis of cognitive aging. Neurobiology of Aging, 33(8), 1699–1715.

Chad, J. A., Pasternak, O., Salat, D. H., & Chen, J. J. (2018). Re-examining age-related differences in white matter microstructure with free-water corrected diffusion tensor imaging. Neurobiology of Aging, 71, 161–170.

Chen, J. J., Rosas, H. D., & Salat, D. H. (2013). The relationship between cortical blood flow and sub-cortical white-matter health across the adult age span. PloS One, 8(2), e56733.

Choy, S. W., Bagarinao, E., Watanabe, H., Ho, E. T. W., Maesawa, S., Mori, D., Hara, K., Kawabata, K., Yoneyama, N., Ohdake, R., Imai, K., Masuda, M., Yokoi, T., Ogura, A., Taoka, T., Koyama, S., Tanabe, H. C., Katsuno, M., Wakabayashi, T., … Sobue, G. (2020). Changes in white matter fiber density and morphology across the adult lifespan: A cross-sectional fixel-based analysis. Human Brain Mapping, 41(12), 3198–3211.

Debette, S., & Markus, H. S. (2010). The clinical importance of white matter hyperintensities on brain magnetic resonance imaging: systematic review and meta-analysis. BMJ, 341, c3666.

de Groot, M., Cremers, L. G. M., Ikram, M. A., Hofman, A., Krestin, G. P., van der Lugt, A., Niessen, W. J., & Vernooij, M. W. (2016). White Matter Degeneration with Aging: Longitudinal Diffusion MR Imaging Analysis. Radiology, 279(2), 532–541.

de Groot, M., Ikram, M. A., Akoudad, S., Krestin, G. P., Hofman, A., van der Lugt, A., Niessen, W. J., & Vernooij, M. W. (2015). Tract-specific white matter degeneration in aging: the Rotterdam Study. Alzheimer’s & Dementia: The Journal of the Alzheimer’s Association, 11(3), 321–330.

Fan, Q., Nummenmaa, A., Witzel, T., Ohringer, N., Tian, Q., Setsompop, K., Klawiter, E. C., Rosen, B. R., Wald, L. L., & Huang, S. Y. (2020). Axon diameter index estimation independent of fiber orientation distribution using high-gradient diffusion MRI. NeuroImage, 222, 117197.

Fan, Q., Tian, Q., Ohringer, N. A., Nummenmaa, A., Witzel, T., Tobyne, S. M., Klawiter, E. C., Mekkaoui, C., Rosen, B. R., Wald, L. L., Salat, D. H., & Huang, S. Y. (2019). Age-related alterations in axonal microstructure in the corpus callosum measured by high-gradient diffusion MRI. NeuroImage, 191, 325–336.

Fjell, A. M., McEvoy, L., Holland, D., Dale, A. M., Walhovd, K. B., & Alzheimer’s Disease Neuroimaging Initiative. (2014). What is normal in normal aging? Effects of aging, amyloid and Alzheimer’s disease on the cerebral cortex and the hippocampus. Progress in Neurobiology, 117, 20–40.

Fjell, A. M., Walhovd, K. B., Fennema-Notestine, C., McEvoy, L. K., Hagler, D. J., Holland, D., Brewer, J. B., & Dale, A. M. (2009). One-year brain atrophy evident in healthy aging. The Journal of Neuroscience: The Official Journal of the Society for Neuroscience, 29(48), 15223–15231.

Grotheer, M., Rosenke, M., Wu, H., Kular, H., Querdasi, F. R., Natu, V. S., Yeatman, J. D., & Grill-Spector, K. (2022). White matter myelination during early infancy is linked to spatial gradients and myelin content at birth. Nature Communications, 13(1), 997.

Han, A., Dhollander, T., Sun, Y. L., Chad, J. A., & Chen, J. J. (2023). Fiber-specific age-related differences in the white matter of healthy adults uncovered by fixel-based analysis. Neurobiology of Aging, 130, 22–29.

Harms, M. P., Somerville, L. H., Ances, B. M., Andersson, J., Barch, D. M., Bastiani, M., Bookheimer, S. Y., Brown, T. B., Buckner, R. L., Burgess, G. C., Coalson, T. S., Chappell, M. A., Dapretto, M., Douaud, G., Fischl, B., Glasser, M. F., Greve, D. N., Hodge, C., Jamison, K. W., … Yacoub, E. (2018). Extending the Human Connectome Project across ages: Imaging protocols for the Lifespan Development and Aging projects. NeuroImage, 183, 972–984.

Hoagey, D. A., Rieck, J. R., Rodrigue, K. M., & Kennedy, K. M. (2019). Joint contributions of cortical morphometry and white matter microstructure in healthy brain aging: A partial least squares correlation analysis. Human Brain Mapping, 40(18), 5315–5329.

Huang, H., Xue, R., Zhang, J., Ren, T., Richards, L. J., Yarowsky, P., Miller, M. I., & Mori, S. (2009). Anatomical characterization of human fetal brain development with diffusion tensor magnetic resonance imaging. The Journal of Neuroscience: The Official Journal of the Society for Neuroscience, 29(13), 4263–4273.

Huang, S. Y., Tian, Q., Fan, Q., Witzel, T., Wichtmann, B., McNab, J. A., Daniel Bireley, J., Machado, N., Klawiter, E. C., Mekkaoui, C., Wald, L. L., & Nummenmaa, A. (2020). High-gradient diffusion MRI reveals distinct estimates of axon diameter index within different white matter tracts in the in vivo human brain. Brain Structure & Function, 225(4), 1277–1291.

Hu, H.-Y., Ou, Y.-N., Shen, X.-N., Qu, Y., Ma, Y.-H., Wang, Z.-T., Dong, Q., Tan, L., & Yu, J.-T. (2021). White matter hyperintensities and risks of cognitive impairment and dementia: A systematic review and meta-analysis of 36 prospective studies. Neuroscience and Biobehavioral Reviews, 120, 16–27.

Kezele, I. B., Arnold, D. L., & Collins, D. L. (2008). Atrophy in white matter fiber tracts in multiple sclerosis is not dependent on tract length or local white matter lesions. Multiple Sclerosis, 14(6), 779–785.

Kiely, M., Triebswetter, C., Cortina, L. E., Gong, Z., Alsameen, M. H., Spencer, R. G., & Bouhrara, M. (2022). Insights into human cerebral white matter maturation and degeneration across the adult lifespan. NeuroImage, 247, 118727.

Kim, C.-M., Alvarado, R. L., Stephens, K., Wey, H.-Y., Wang, D. J. J., Leritz, E. C., & Salat, D. H. (2020). Associations between cerebral blood flow and structural and functional brain imaging measures in individuals with neuropsychologically defined mild cognitive impairment. Neurobiology of Aging, 86, 64–74.

Kodiweera, C., Alexander, A. L., Harezlak, J., McAllister, T. W., & Wu, Y.-C. (2016). Age effects and sex differences in human brain white matter of young to middle-aged adults: A DTI, NODDI, and q-space study. NeuroImage, 128, 180–192.

Liewald, D., Miller, R., Logothetis, N., Wagner, H.-J., & Schüz, A. (2014). Distribution of axon diameters in cortical white matter: an electron-microscopic study on three human brains and a macaque. Biological Cybernetics, 108(5), 541–557.

Madden, D. J., Bennett, I. J., Burzynska, A., Potter, G. G., Chen, N.-K., & Song, A. W. (2012). Diffusion tensor imaging of cerebral white matter integrity in cognitive aging. Biochimica et Biophysica Acta, 1822(3), 386–400.

Maffei, C., Lee, C., Planich, M., Ramprasad, M., Ravi, N., Trainor, D., Urban, Z., Kim, M., Jones, R. J., Henin, A., Hofmann, S. G., Pizzagalli, D. A., Auerbach, R. P., Gabrieli, J. D. E., Whitfield-Gabrieli, S., Greve, D. N., Haber, S. N., & Yendiki, A. (2021). Using diffusion MRI data acquired with ultra-high gradients to improve tractography in routine-quality data. In bioRxiv (p. 2021.06.28.450265). 10.1101/2021.06.28.450265

Marner, L., Nyengaard, J. R., Tang, Y., & Pakkenberg, B. (2003). Marked loss of myelinated nerve fibers in the human brain with age. The Journal of Comparative Neurology, 462(2), 144–152.

Mosconi, L., Berti, V., Dyke, J., Schelbaum, E., Jett, S., Loughlin, L., Jang, G., Rahman, A., Hristov, H., Pahlajani, S., Andrews, R., Matthews, D., Etingin, O., Ganzer, C., de Leon, M., Isaacson, R., & Brinton, R. D. (2021). Menopause impacts human brain structure, connectivity, energy metabolism, and amyloid-beta deposition. Scientific Reports, 11(1), 10867.

Ouyang, Y., Cui, D., Yuan, Z., Liu, Z., Jiao, Q., Yin, T., & Qiu, J. (2021). Analysis of Age-Related White Matter Microstructures Based on Diffusion Tensor Imaging. Frontiers in Aging Neuroscience, 13, 664911.

Pakkenberg, B., & Gundersen, H. J. (1997). Neocortical neuron number in humans: effect of sex and age. The Journal of Comparative Neurology, 384(2), 312–320.

Piguet, O., Double, K. L., Kril, J. J., Harasty, J., Macdonald, V., McRitchie, D. A., & Halliday, G. M. (2009). White matter loss in healthy ageing: a postmortem analysis. Neurobiology of Aging, 30(8), 1288–1295.

Sala, S., Agosta, F., Pagani, E., Copetti, M., Comi, G., & Filippi, M. (2012). Microstructural changes and atrophy in brain white matter tracts with aging. Neurobiology of Aging, 33(3), 488–498.e2.

Schilling, K. G., Archer, D., Yeh, F.-C., Rheault, F., Cai, L. Y., Hansen, C., Yang, Q., Ramdass, K., Shafer, A. T., Resnick, S. M., Pechman, K. R., Gifford, K. A., Hohman, T. J., Jefferson, A., Anderson, A. W., Kang, H., & Landman, B. A. (2022). Aging and white matter microstructure and macrostructure: a longitudinal multi-site diffusion MRI study of 1218 participants. Brain Structure & Function, 227(6), 2111–2125.

Seiler, S., Fletcher, E., Hassan-Ali, K., Weinstein, M., Beiser, A., Himali, J. J., Satizabal, C. L., Seshadri, S., DeCarli, C., & Maillard, P. (2018). Cerebral tract integrity relates to white matter hyperintensities, cortex volume, and cognition. Neurobiology of Aging, 72, 14–22.

Stadlbauer, A., Salomonowitz, E., Strunk, G., Hammen, T., & Ganslandt, O. (2008). Age-related degradation in the central nervous system: assessment with diffusion-tensor imaging and quantitative fiber tracking. Radiology, 247(1), 179–188.

Taha, H. T., Chad, J. A., & Chen, J. J. (2022). DKI enhances the sensitivity and interpretability of age-related DTI patterns in the white matter of UK biobank participants. Neurobiology of Aging, 115, 39–49.

Taki, Y., Kinomura, S., Sato, K., Goto, R., Wu, K., Kawashima, R., & Fukuda, H. (2011). Correlation between gray/white matter volume and cognition in healthy elderly people. Brain and Cognition, 75(2), 170–176.

Tang, Y., Nyengaard, J. R., Pakkenberg, B., & Gundersen, H. J. (1997). Age-induced white matter changes in the human brain: a stereological investigation. Neurobiology of Aging, 18(6), 609–615.

Uddin, M. N., Figley, T. D., Solar, K. G., Shatil, A. S., & Figley, C. R. (2019). Comparisons between multi-component myelin water fraction, T1w/T2w ratio, and diffusion tensor imaging measures in healthy human brain structures. Scientific Reports, 9(1), 2500.

Wu, H., Sun, C., Huang, X., Wei, R., Li, Z., Ke, D., Bai, R., & Liang, H. (2022). Short-Range Structural Connections Are More Severely Damaged in Early-Stage MS. AJNR. American Journal of Neuroradiology, 43(3), 361–367.

Yeatman, J. D., Wandell, B. A., & Mezer, A. A. (2014). Lifespan maturation and degeneration of human brain white matter. Nature Communications, 5, 4932.

Yendiki, A., Panneck, P., Srinivasan, P., Stevens, A., Zöllei, L., Augustinack, J., Wang, R., Salat, D., Ehrlich, S., Behrens, T., Jbabdi, S., Gollub, R., & Fischl, B. (2011). Automated probabilistic reconstruction of white-matter pathways in health and disease using an atlas of the underlying anatomy. Frontiers in Neuroinformatics, 5, 23.

